# Prevalence of sustainable and unsustainable use of wild species inferred from the IUCN Red List

**DOI:** 10.1101/2020.11.04.367763

**Authors:** Sophie M.E. Marsh, Michael Hoffmann, Neil D. Burgess, Thomas M. Brooks, Daniel W.S. Challender, Patricia J. Cremona, Craig Hilton-Taylor, Flore Lafaye de Micheaux, Gabriela Lichtenstein, Dilys Roe, Monika Böhm

**Affiliations:** Centre for Biodiversity and Environment Research, Department of Genetics, Evolution and Environment, University College London, Gower Street, London WC1E 6BT, UK.; Conservation and Policy, Zoological Society of London, Regent’s Park, London, NW1 4RY.; UNEP-WCMC, 219 Huntington Road, CB3 0DL, Cambridge, UK; CMEC, GLOBE Institute, University of Copenhagen, Denmark; International Union for Conservation of Nature, Gland, Switzerland; World Agroforestry Center (ICRAF), University of the Philippines, Los Baños, The Philippines; Institute for Marine and Antarctic Studies, University of Tasmania, Hobart, Australia; Department of Zoology, University of Oxford, Zoology Research and Administration Building, 11a Mansfield Road, Oxford, OX1 3SZ, United Kingdom; International Union for Conservation of Nature, Cambridge, UK; Institute of Geography and Sustainability, University of Lausanne, Switzerland; French Institute of Pondicherry, India; Instituto Nacional de Antropología y Pensamiento Latinoamericano (INAPL)/CONICET, Argentina; International Institute for Environment and Development (IIED) and IUCN Sustainable Use and Livelihoods Specialist Group (SULi), London UK; Institute of Zoology, Zoological Society of London, Regent’s Park, London, NW1 4RY

**Keywords:** Policy, wildlife, (un)sustainable use, exploitation, IPBES, Convention on Biological Diversity, CITES, conservation action

## Abstract

Unsustainable exploitation of wild species represents a serious threat to biodiversity and to the livelihoods of local communities and indigenous peoples. However, managed, sustainable use has the potential to forestall extinctions, aid recovery, and meet human needs. Research to date has focused on unsustainable biological resource use with little consideration of sustainable use; we infer the current prevalence of both. We analyzed species-level data for 30,923 species from 13 taxonomic groups on the IUCN Red List. Our results demonstrate the broad taxonomic prevalence of use, with 40% of species (10,098 of 25,009 species from 10 data-sufficient taxonomic groups) documented as being used. The main purposes of use are pets, display animals and horticulture, and human consumption. Use often has an adverse impact on species extinction risk (we define this as biologically unsustainable): intentional use is currently contributing to elevated extinction risk for over a quarter of all threatened or Near Threatened (NT) species (2,752 – 2,848 of 9,753 species). Intentional use also threatens 16% of all species used (1,597 – 1,631 of 10,098). However, 72% of species that are used (7,291 of 10,098) are Least Concern (LC), of which nearly half (3,469) also have stable or improving population trends. The remainder of used species are not documented as threatened by biological resource use, including 172 threatened or NT species with stable or improving populations. Around a third of species that have use documented as a threat do not currently receive targeted species management actions to directly address this threat. We offer suggestions for improving use-related Red List data. Our findings on the prevalence of sustainable and unsustainable use, and variation across taxa, can inform international policymaking, including the Intergovernmental Science-Policy Platform on Biodiversity and Ecosystem Services, the Convention on Biological Diversity, and the Convention on International Trade in Endangered Species.

## Introduction

It is critical to understand and manage the impacts of threats related to the use of wild species to ensure their survival while continuing to support global demand for biological resources. Over-exploitation is among the predominant threats to many species (Maxwell et al., 2016; di Minin et al. 2019), and the primary threat to aquatic species (IPBES 2019). Nonetheless, billions of people rely on wild species, including plants, animals and fungi, for their food, medicines, construction and other uses (Nasi et al. 2011; Thilsted et al. 2016). Indeed, the use of wild species underpins the livelihoods of millions of people and has cultural, religious and recreational value. These values in turn provide a local incentive for the conservation of species. The tension between over-exploitation as a major driver of biodiversity loss, and humanity’s reliance on wild species for many different needs creates a conundrum. How can biological resource use be managed in sustainable ways that help meet human needs and incentivize conservation, rather than further driving species to extinction?

The use of wild species can be sustainable given adequate management (Lichtenstein 2010; Austin & Corey 2012). The concept of sustainable use is embedded in many international and national regulatory and policy frameworks as a conservation management tool, to promote human development, and to ensure availability of natural resources for future generations. It is one of the three primary objectives of the Convention on Biological Diversity (CBD), and is explicitly stated in the UN Sustainable Development Goals. Nonetheless, sustainable use as a practice remains polarizing (Hutton & Leader-Williams 2003; Challender & MacMillan 2019), especially consumptive use of animals (involving the removal of either live or dead individuals), and with limited consensus regarding the effectiveness of different approaches. This issue is exacerbated by concerns that inaction or ineffective sustainable use policies could rapidly imperil many already threatened species (Auliya et al. 2016). Yet conversely, actions to prevent or reduce use could have negative consequences (Cooney & Jepson 2006; Bonwitt et al. 2018), for species concerned and for people who depend on their use, in particular those most vulnerable.

The discourse around sustainable use is further hampered by knowledge gaps. While there is a good body of research around how use and trade drive declines and imperil species, we have limited understanding of different patterns of use within species. The same can be said for the degree to which species currently being impacted by over-exploitation are receiving appropriate conservation actions, or where, and under what circumstances, trade is taking place at sustainable levels without threatening species (Morton et al. 2021). Several data sources which can provide insights into sustainable versus unsustainable use exist, but may be limited in geographical or taxonomic scope. For example, McRae et al. (2021) used a global dataset of more than 11,000 population time series from the Living Planet Database to derive trends in utilized versus non-utilized vertebrates, and to assess whether ‘management’ makes a measurable difference to wildlife population trends for utilized species. While the underlying data are population-specific and provide a higher resolution than species-level databases, the taxonomic scope is limited to vertebrates only. Other datasets can provide taxon- and region-specific insight into the use of species, including Prota4U for plants (https://www.prota4u.org/) and national Red Lists. However, they are not directly comparable for global analyses, not least because they use different taxonomies and have different protocols for capturing information on use.

The IUCN Red List of Threatened Species (henceforth ‘Red List’) provides global data for a wide range of taxa that can assist managers and policymakers in understanding and delivering targeted action to address threats to biodiversity. The role of the Red List in supporting and influencing global policy instruments is well established, for instance in tracking progress towards globally agreed targets such as the CBD Aichi Targets (SCBD 2014), new targets under discussion in the post-2020 Global Biodiversity Framework, and Sustainable Development Goals (Brooks et al. 2015). The Red List also provides key data and trends that inform processes in the Convention on International Trade in Endangered Species of Fauna and Flora (CITES; Challender et al. 2019) and the Intergovernmental Science-Policy Platform on Biodiversity and Ecosystem Services (IPBES; Brooks et al. 2016; IPBES 2019).

Individual Red List assessments are carried out by thousands of scientific experts in accordance with a system of objective, quantitative categories and criteria that rank a species’ extinction risk from Least Concern (LC) to Extinct in the Wild (EW), or Extinct (EX). A species is considered threatened if it is assessed as Vulnerable (VU), Endangered (EN), or Critically Endangered (CR) (IUCN 2012). Assessments follow well-defined guidelines with an independent process for review (Collen et al. 2016), and are underpinned by ancillary data on distribution, population size and trend, habitat preferences, threats and conservation actions in place or needed. Much of this information is recorded in standardized “Classification Schemes” that enable comparative analyses across taxa.

Previous analyses of biological resource use using Red List data have focused on individual taxonomic groups (e.g. birds, Butchart 2008; cacti, Goettsch et al. 2015), or on particular dimensions of use (e.g. traded vertebrates, Scheffers et al. 2019; spatial concentrations of unsustainable use, di Minin et al. 2019). We substantially advance previous analyses of Red List data by investigating all patterns of intentional biological resource use across a broad suite of species assessed on the Red List. We ask: 1) what are the main purposes of use of wild animal and plant species; 2) for which species are current levels of use having a negative impact on species extinction risk, and hence likely biologically unsustainable; 3) for which species are current levels of use not having a negative impact on species extinction risk, and hence likely biologically sustainable; and 4) for which utilized species are conservation actions currently in place to directly target impacts from current levels of use. Our study provides a framework for replicating the results in the future (for example, to track trends over time), and yields concrete suggestions for improving the quality of use-related Red List data in future assessments.

## Methods

### Species data

We collated species-level data for 13 taxonomic groups that have been comprehensively assessed on the Red List (version 2020-1). The Red List defines comprehensively assessed groups as taxonomic groups that include at least 150 species, of which >80% have been assessed (IUCN 2020). Non-comprehensively assessed groups may primarily focus on species that are likely threatened or likely used by people, or may have a regional focus. Excluding these groups avoids introducing biases into our analysis, as for instance threat processes are usually not evenly distributed geographically (Miqueleiz et al. 2020).

We classed the 13 taxonomic groups into six primarily aquatic (freshwater and/or marine) and seven primarily terrestrial groups (including amphibians, among which ∼30% are documented as terrestrial only, the remainder as both terrestrial and freshwater). We excluded all species listed as Extinct (EX) or Extinct in the Wild (EW), as neither can be currently used in the wild, or Data Deficient (DD), as the impact of any use on their extinction risk is unknown. This restricted our analyses to Least Concern (LC), Near Threatened (NT), and threatened species only (hereafter ‘extant, data sufficient species’). This dataset comprises 30,923 species, made up of 6,603 primarily aquatic species and 24,320 primarily terrestrial species (Table 1; Supporting Information).

**Table 1.**
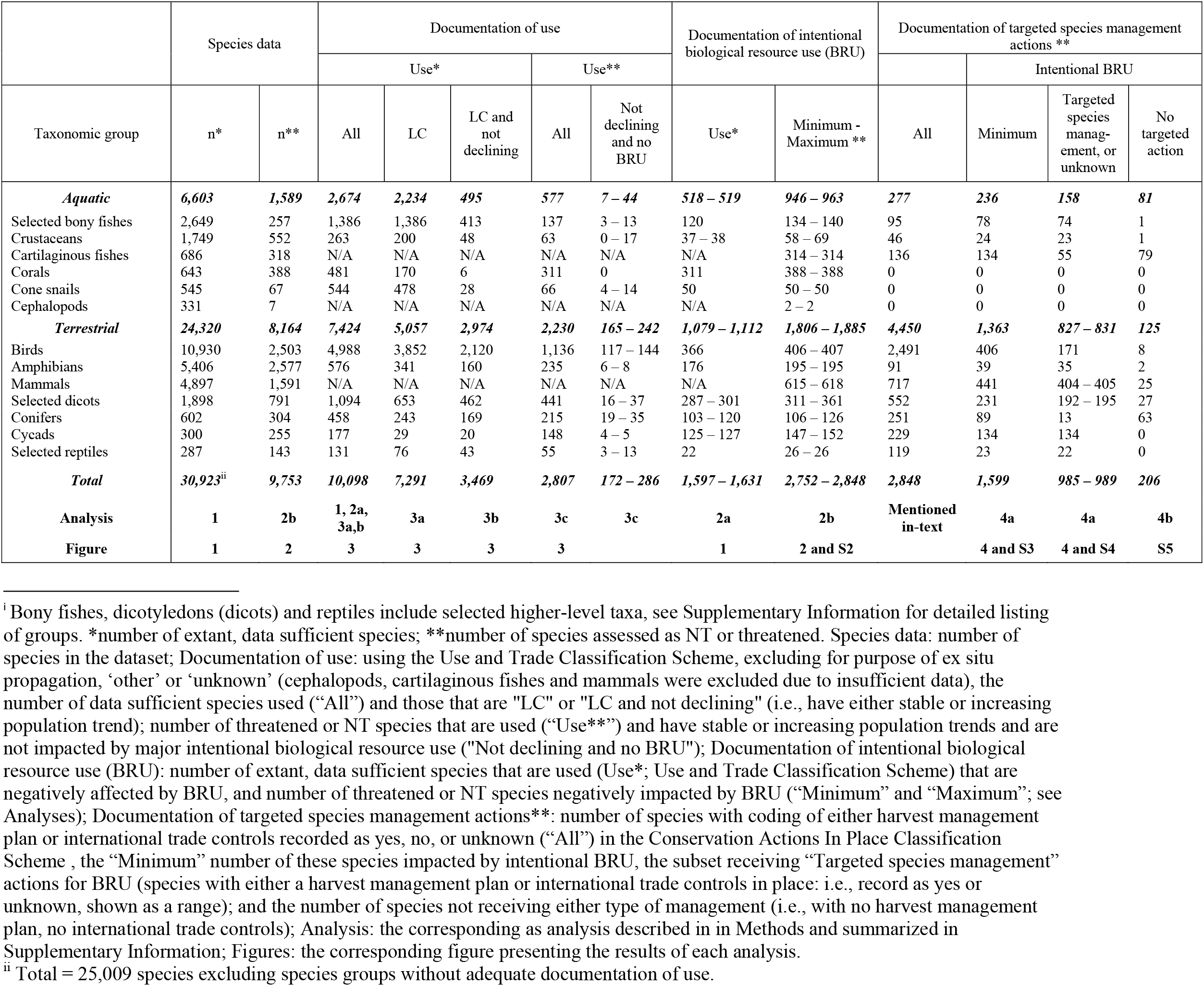
Comprehensively assessed species groups and subsets included in analyses and their respective sample sizes.^i^

Note that our dataset of extant, data sufficient species includes 12% which have not been (re)assessed in the last 10 years (this is 22% of those threatened or NT). This proportion is lower than for the overall population of species with outdated assessments (version 2020-3) on the Red List (23.8%). We acknowledge that these older assessments may not account properly for current patterns of use (equally, their extinction risk categories, threats, current population trends and other supporting information might have changed). However, we consider it appropriate to include these species since these data remain the best we have, and were compiled as part of major global assessment processes that did include explicit documentation of use. Complete species lists for all analyses are available from the authors.

### Red List data used

For each species, we downloaded the following data from the Red List: Red List category, current population trend, threats, use and trade, and conservation actions in place. Information on the latter three is recorded for each species using the Red List’s standardized Classification Schemes, which provide a harmonized typology for recording relevant attributes (Salafsky et al. 2008). Documenting current population trends – where current reflects the time of assessment – is mandatory to publication of all species assessments on the Red List and is presented as either stable, decreasing, increasing, or unknown. Use of species is captured on the Red List in two distinct ways: as a threat under the Threats Classification Scheme (Class 5, Biological Resource Use; Salafsky et al. 2008; Supporting Information), and as a form of use or trade under the Use and Trade Classification Scheme, which explicitly does not associate the use with a threat (Supporting Information). The threat information can tell us whether species are negatively impacted by use, whereas the use and trade information documents the purposes of use regardless of whether it represents a threat or not. While the recording of major threats impacting a species is mandatory documentation for species listed as EX, EW, threatened, and NT (IUCN 2016), recording of use and trade is only recommended (i.e. strongly encouraged, but not mandatory for publication) documentation and may thus not be consistently documented across all species on the Red List. Conservation actions are recorded in the Conservation Actions In Place Classification Scheme and Conservation Actions Needed Classification Scheme. In both schemes, this information is also only recommended documentation, and thus may not be consistently recorded (Luther et al. 2016). For the purpose of this study, we only analyzed the Conservation Actions In Place Classification Scheme.

### Analyses

#### The main purposes of use of wild animal and plant species on the Red List

We investigated the prevalence of different purposes of use based on the information recorded in the Use and Trade Classification Scheme, excluding records for “establishing ex situ production” (use code 16), “other” (17), and “unknown” (18). For this analysis, we limited our dataset to 10 taxonomic groups that have adequate recording of use and trade, leading us to exclude cartilaginous fishes and cephalopods from the aquatic species group, and mammals (a high-profile group when it comes to discussion of use) from the terrestrial group. For each taxonomic group, we calculated the total number of species recorded as being used for at least one purpose in the Use and Trade Classification Scheme. We then summarized results as the percentage of species recorded for different types of use on the Red List out of all extant, data sufficient species (totaling 25,009 species in these 10 taxonomic groups; Table 1). A detailed description of how adequate recording of use was defined and the full list of selected groups can be found in Supporting Information.

#### Wild species for which intentional use is having a negative impact on extinction risk

For our analysis, we considered use to be biologically unsustainable when it is likely to be having an adverse impact on species extinction risk. To identify such cases, we analyzed the proportion of species for which “biological resource use” is documented as a major threat, as recorded using the IUCN Threat Classification Scheme, from among: a) all species with at least one purpose of use coded (from among the 10 taxonomic groups with adequate data, see previous section on main purposes of use), and b) all NT and threatened species (from all 13 taxonomic groups comprehensively assessed) (Table 1). Since not all types of biological resource use are intentionally targeting the species in question, we developed a decision-tree for removing threat types that are not relevant to an analysis of direct, intentional use of species (Supporting Information). In some cases, we were unable to determine whether the use was intentional. We present this uncertainty in our results as a range where the “minimum” proportion includes all species with threats that could be conclusively determined as intentional. The “maximum” proportion additionally includes species with motivation unknown or unrecorded that may represent further cases of intentional biological resource use. For groups where no species have such motivation unknown / unrecorded threats, we present only the minimum (Supporting Information).

We included biological resource use as a threat if it had a medium to high impact on species extinction risk. The Red List uses a scoring system to estimate threat impact, based on timing, scope, and severity of the threat. This information is used to create an overall threat impact score. Threat timing is recommended information to be provided in the Red List assessment (i.e. strongly encouraged but not essential to publication), while severity and scope are only discretionary and often not included. As such, we amended the threat impact score categorization to help exclude threats that are not likely to be major (Supporting Information).

#### Wild species for which intentional use is not having a negative impact on extinction risk

We considered use to be likely biologically sustainable when it is unlikely to have an adverse impact on species extinction risk. Due to data constraints, we could only derive a minimum estimate by determining the number of species recorded as being subject to some form of use or trade that: a) are currently LC, b) LC and not declining (i.e., have either stable or increasing current population trends); and c) are threatened or NT, not documented with intentional use as a major threat, and have stable or increasing population trends (Table 1). We recognize the limitations to using overall current population trends in this analysis, but we consider this to be defensible. Our first approach ties population trend to extinction risk (specifically, if a species is LC then intentional use or trade, or indeed any threat, cannot, by definition, be a major threat), while our second approach is explicitly tied to threatened or NT species that do not have use or trade documented as a major threat, and as such this cannot be a primary factor driving elevated extinction risk. We confined analyses to those 10 taxonomic groups for which use and trade information was adequate, as discussed above.

#### Conservation actions in place or lacking for utilized wild species

To understand the current level of conservation actions in place to respond to over-exploitation, we extracted all NT and threatened species that are documented as receiving targeted species management actions, as recorded via the Conservation Actions In Place Classification Scheme. Specifically, we selected species for which there is a harvest management plan in place and species subject to any international management / trade controls (e.g., CITES or US Endangered Species Act listing, regional fisheries agreements). We then determined the number of species that are negatively impacted by biological resource use, and: a) are recorded as receiving either one or both of these actions, and b) are not receiving either of these actions (Table 1).

## Results

### The main purposes of use of wild animal and plant species on the Red List

Among the 10 taxonomic groups with adequate information, the proportion of extant, data sufficient species documented as having at least one purpose of use coded ranged from 15% (crustaceans) to nearly 100% of cone snails (544 of 545 species) among aquatic groups, and 11% (amphibians) to 76% (conifers) among terrestrial groups. Across the 25,009 species in these 10 groups, 10,098 (40%) had some purpose of use documented (Table 1).

In the aquatic groups, the top purposes of use were for human food (selected bony fishes and crustaceans), specimen collection (cone snails), and pets and display animals (corals and selected bony fishes) (Fig. 1). Additional purposes of use were for handicrafts and jewelry (cone snails and corals) and medical purposes (cone snails). For terrestrial animal groups, the two most prevalent uses were for pets or display animals and for human consumption (birds and herptiles). This was followed by sport hunting and specimen collecting for birds, medicinal purposes for amphibians and wearing apparel / accessories for selected reptiles. For plant taxonomic groups, the predominant uses were for structural and building materials (conifers) and horticulture (all three groups). Overall, plant groups were used for more purposes than animal groups, including for human and animal food, medicinal use, household goods and handicrafts / jewelry, fuels and chemicals.

**Figure 1.**
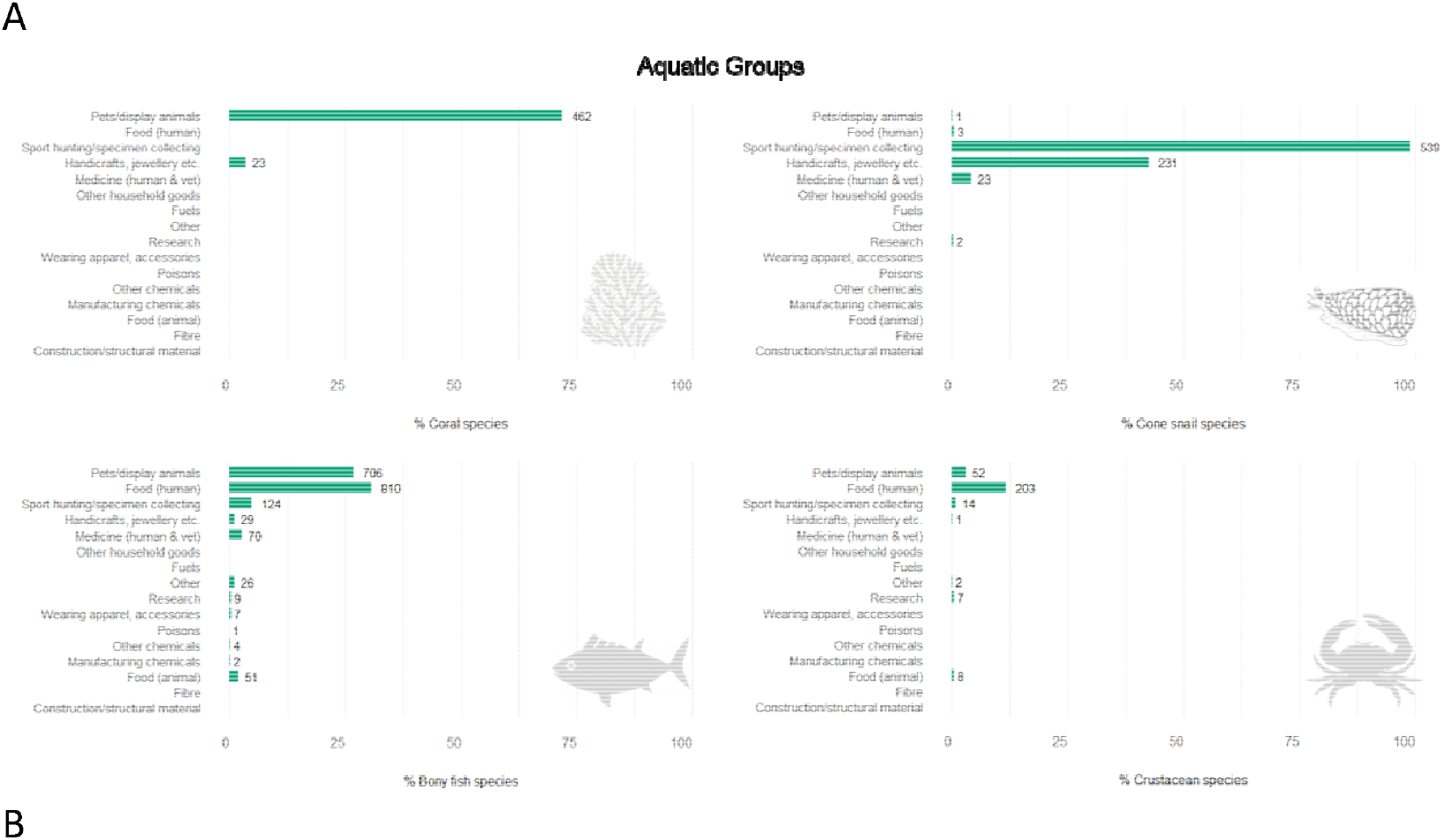
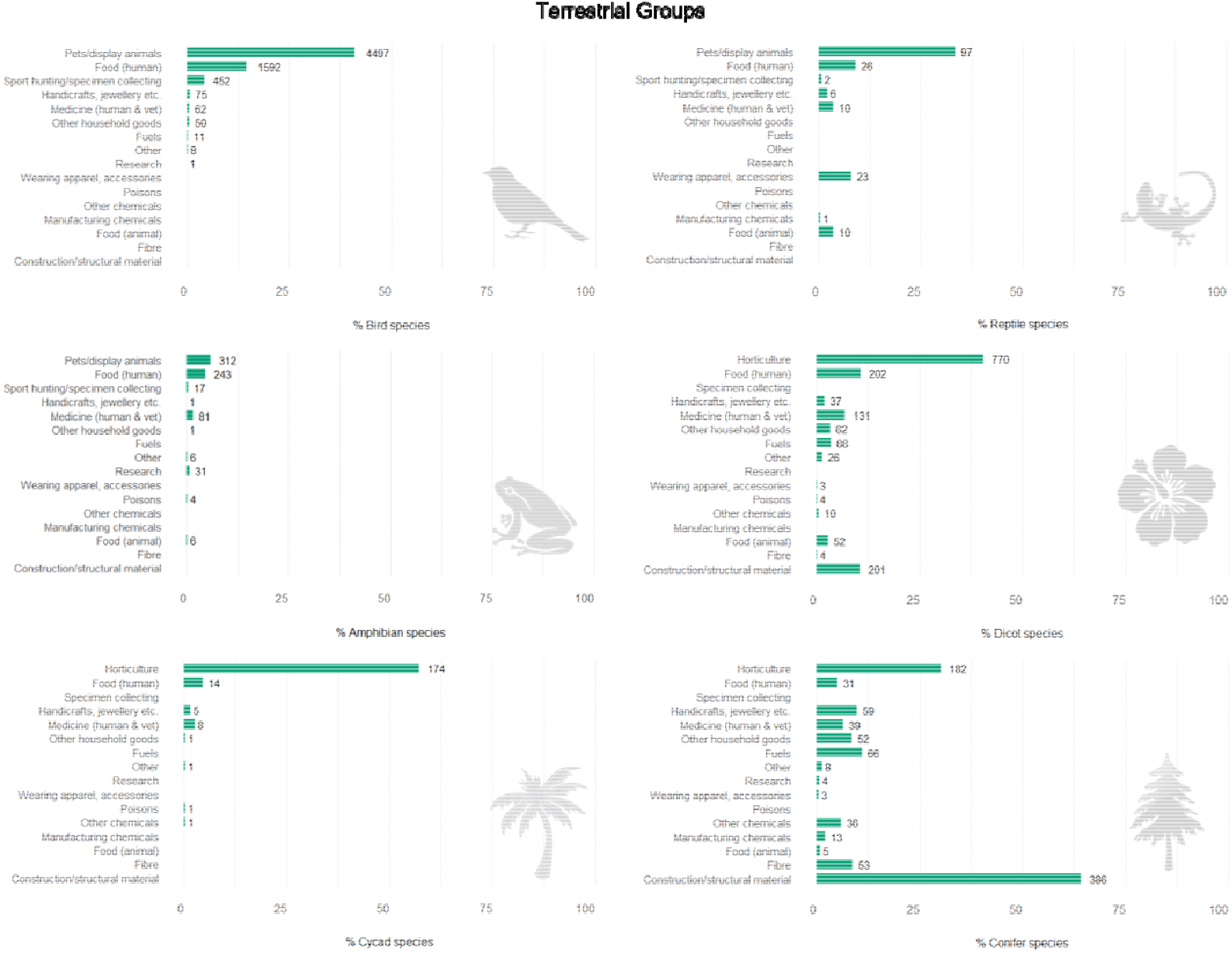
Percentage of extant, data sufficient species in A) aquatic and B) terrestrial taxonomic groups, recorded for different types of use on the Red List. Percentages out of total extant, data sufficient species (see Table 1). Data labels show the total number of species recorded for each purpose of use. Note that most species are subject to more than one type of use. Bony fishes, dicotyledons (dicots) and reptiles include selected higher-level taxa (Supplementary Information).

#### Wild species for which intentional use is having a negative impact on extinction ris

Considering all 10,098 species for which some purpose of use is documented in the 10 taxonomic groups with adequate information, a sixth have intentional biological resource use documented as a threat (1,597 – 1,631 species, including those with motivation unknown, representing 16%). Moreover, more than a quarter of all NT and threatened species, across all 13 comprehensively assessed taxa (9,753 species), have intentional biological resource use documented as a threat (minimum = 2,752 species, or 28%; maximum = 2,848, or 29%).

Across NT and threatened species, a higher overall proportion of aquatic species than terrestrial species have intentional biological resource use recorded as a threat (Fig. 2). Among aquatic groups, the taxa with highest prevalence are corals (100%; 388 species) and almost all cartilaginous fishes (99%; 314 out of 318 species), with fishing and harvesting the predominant threat (Supporting Information); in the terrestrial groups, cycads appear most impacted (58 – 60%, 147 – 152 of 255 species), largely due to collecting (147 species).

**Figure 2.**
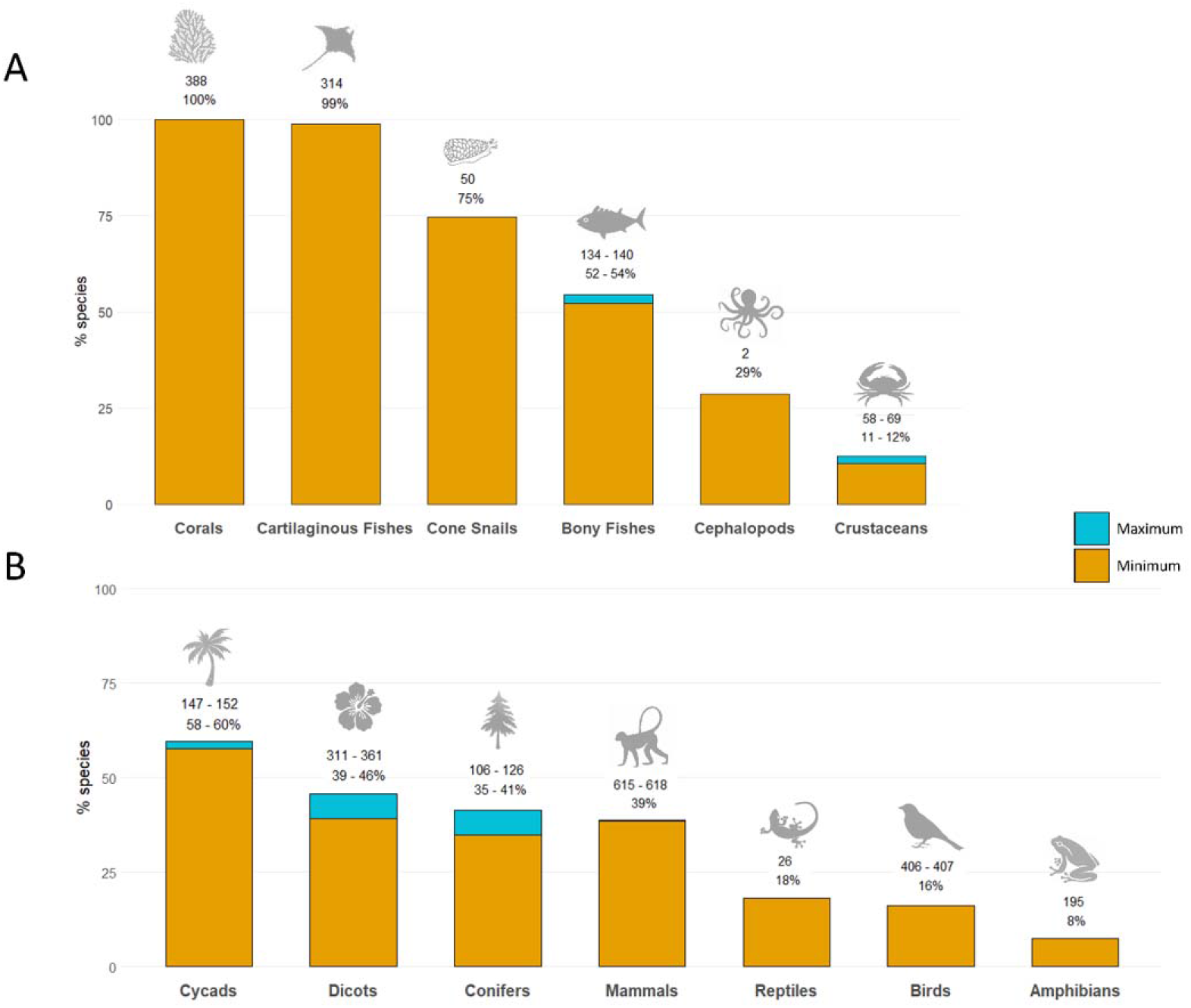
Percentage of NT and threatened species in A) aquatic and B) terrestrial groups with biological resource use documented as a threat on the Red List. Minimum (orange) bars are defined by the number of species in each taxonomic group that are affected by at least one type of intentional use; maximum (blue) bars are species that might be subject to intentional use, including where species are coded as affected by use under “motivation unknown”. Black labels denote the minimum number of species affected by biological resource use in each taxonomic group, and the percentage range from minimum to maximum (where relevant). Bony fishes, dicotyledons (dicots) and reptiles include selected higher-level taxa (Supplementary Information).

#### Wild species for which intentional use is not having a negative impact on extinction risk

Among the 10 taxonomic groups for which information was adequate, most species subjected to some form of use or trade were LC, with the exception of cycads and corals. The overall percentage of utilized species that were LC was 72%, ranging from 16% and 35% in cycads and corals, respectively, to 76% in crustaceans, 77% in birds, 88% in cone snails and 90% in selected bony fishes (Fig. 3). Among terrestrial groups, between 11% (cycads, 20 species) and 42% (birds and selected dicots, 2,120 and 462 species, respectively) of utilized species are LC and also have either stable or increasing population trends (are not currently declining). For aquatic groups, proportions are lower, ranging from 1% (corals, 6 species) to 30% (selected bony fishes, 413 species). Across all 10 taxa for which data on purpose of use are adequate, 34% (3,469 of 10,098) of utilized species are LC and also not currently declining. Furthermore, we documented 172 threatened and NT species (2% of utilized species) subject to some form of use that are not impacted by intentional biological resource use and further exhibit stable or increasing population trends (Table 1).

**Figure 3.**
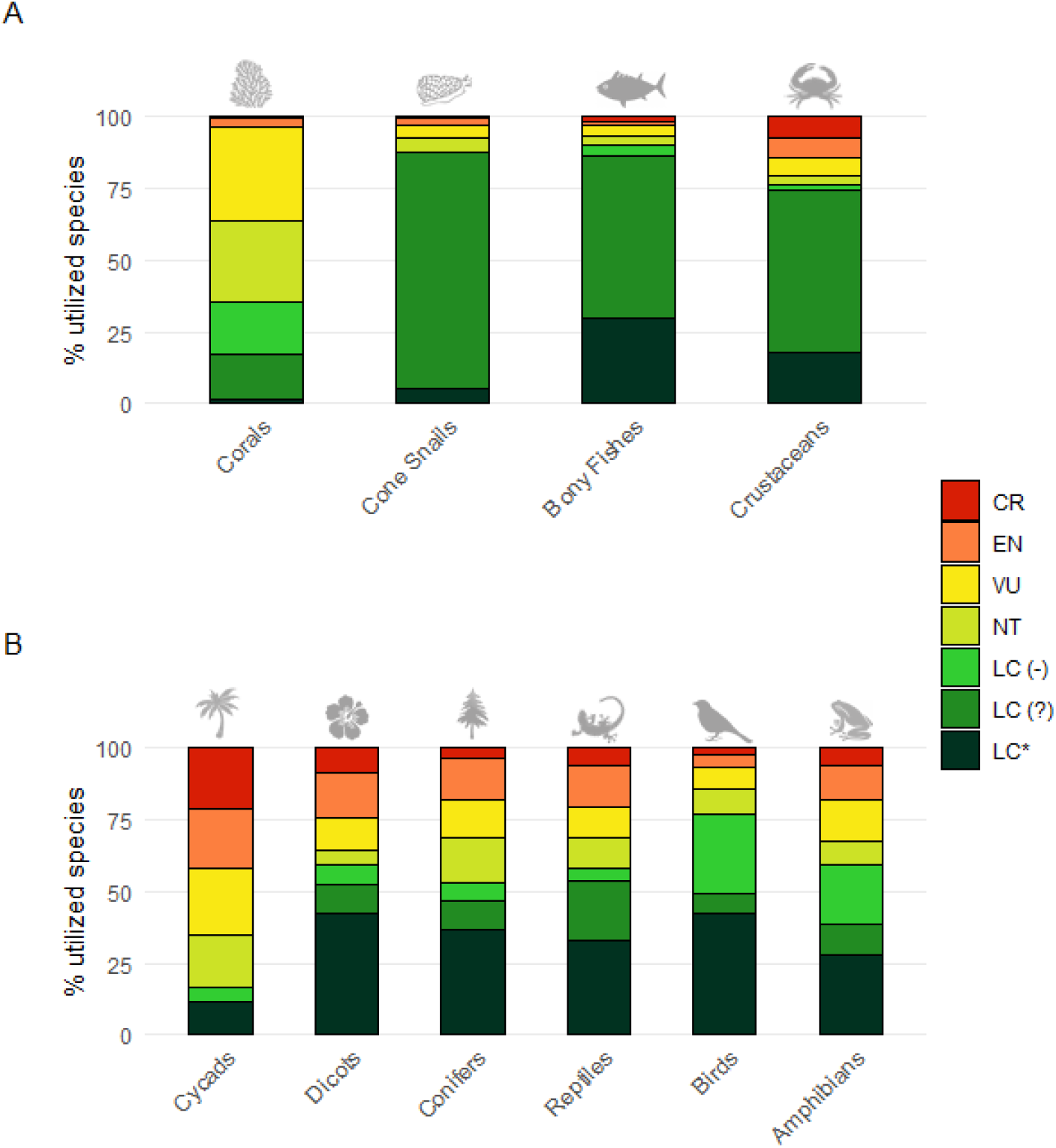
Percentage of extant, data sufficient species by Red List Category in A) aquatic and B) terrestrial groups that are subject to use and trade. LC(-) = Least Concern species with declining population trend; LC(?) = Least Concern species with unknown population trend; LC(*) = Least Concern species with stable or increasing population trend. Note that being LC and having a declining population trend, or being threatened and being subject to use and trade, does not imply that use is a major threat; we have no evidence as to whether use is sustainable or not for 48% of used species. Bony fishes, dicotyledons (dicots) and reptiles include selected higher-level taxa (Supplementary Information).

#### Conservation actions in place or lacking for utilized wild species

No information on species management actions relevant to use is recorded for NT or threatened corals, cone snails or cephalopods (Table 1). Among NT and threatened species impacted by biological resource use, only 1% of cycads (1 species), 3% of birds (12 species), and 6% of amphibians (11 species) have available information on harvest management actions (Supporting Information). There is only recorded data on international trade controls for 3% of crustaceans (2 species) and 6% of amphibians (11 species). However, over 80% of conifers, selected reptiles, and cycads and 100% of birds have available documentation of species harvest management interventions (Supporting Information). From those species for which there is available data for one or both actions, cycads, selected reptiles, mammals and selected dicots all have >80% of species documented as being subject to some form of international trade control, and >80% of crustaceans are subject to harvest management actions (Supporting Information). Species groups are more likely to receive international trade controls (64% of species) than harvest management actions (9%), although conifers (11% and 7%, respectively), selected bony fishes (42% and 59%), and cartilaginous fishes (21% and 24%) receive international trade control measures and harvest management interventions in nearly equal proportions (Fig. 4).

**Figure 4.**
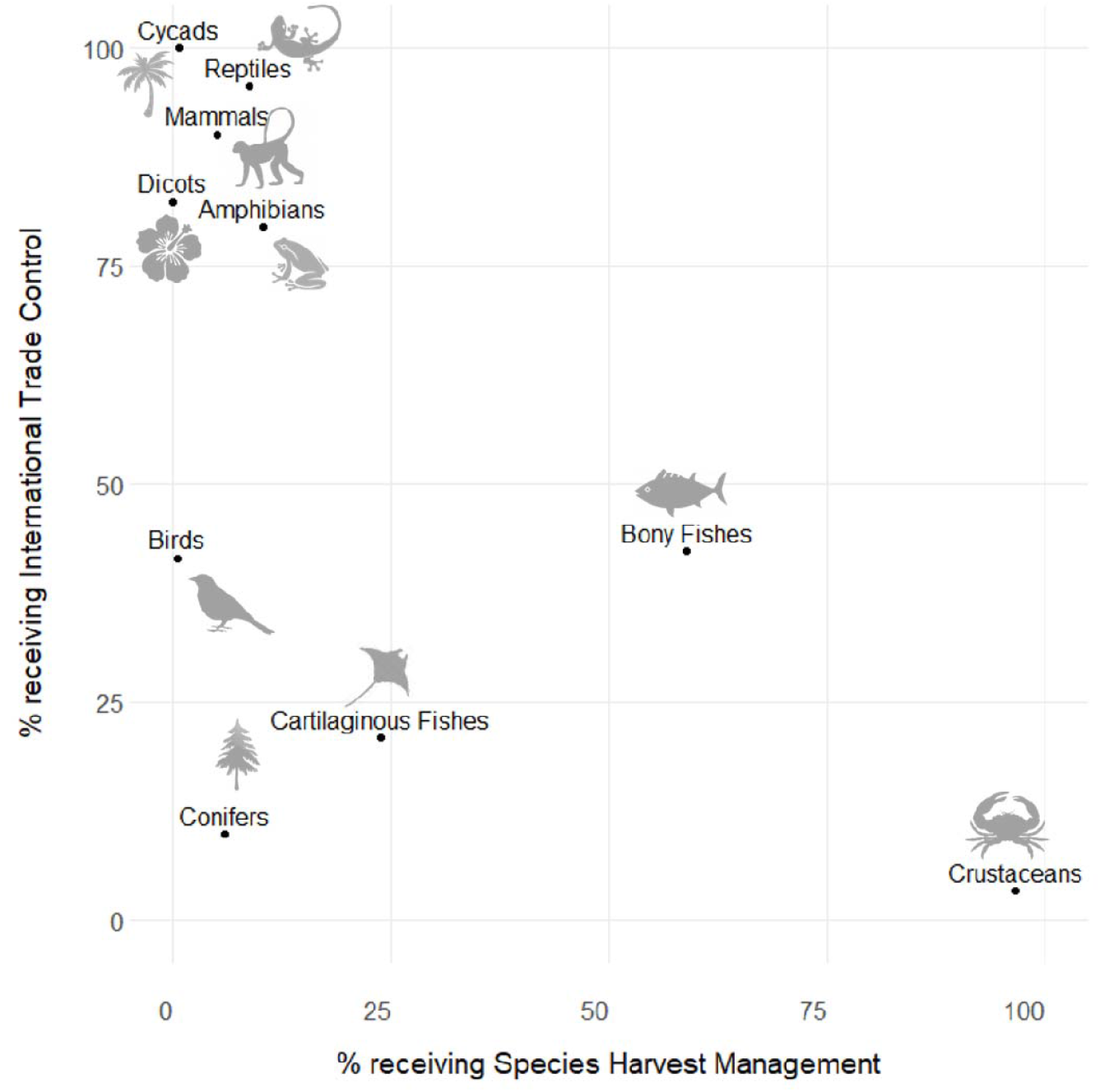
Relationship between the prevalence of international trade controls and species harvest management across NT and threatened species affected by intentional biological resource use (minimum estimate), based on species with available data on conservation actions (those where the field is recorded as either “Unknown,” “Yes,” or “No”, rather than left blank). No data are available for cephalopods, cone snails or corals (see Figure S3). Bony fishes, dicotyledons (dicots) and reptiles include selected higher-level taxa (Supplementary Information).

In total, out of at least 2,752 threatened and NT species that have intentional biological resource use coded as a threat (1,599 of which have available documentation for one or both conservation actions), fewer than a thousand (985 – 989) are documented as benefitting from either international trade control or species harvest management interventions; at least 206 species are explicitly stated as lacking any such actions (Supporting Information). Compared with terrestrial groups (9% of species with available documentation), species in aquatic groups are more likely (34%) to be lacking any conservation management in response to biological resource use (Table 1).

## Discussion

While previous analyses of Red List data have mostly examined the degree to which biological resource use is a threatening process driving extinction risk (which we further expand on here), our study provides a first attempt to analyze Red List data to understand the extent to which use of wild species is *not* having a detrimental impact on species extinction risk. Although our analyses were hindered by data constraints, our findings that nearly three-quarters of species that are used are currently categorized as Least Concern indicate that, at least at the time of assessment, use is not placing these species at risk of extinction. Further, more than one-third each of birds, selected reptiles, conifers and selected dicots that are used are categorized as Least Concern and also exhibit either stable or increasing population trends. We also find evidence that 172 threatened and Near Threatened species used have stable or increasing population trends and do not have biological resource use as a major threat. Red List assessments concern species across their range, consequently it is possible that, despite overall current population trends at the time of assessment, some species could be undergoing localized declines now (or have undergone these in the past) due to the impacts of biological resource use (or indeed localized increases due to successful interventions). There could also be localized increases due to successful interventions. For example, the Northern Chamois (*Rupicapra rupicapra*) is currently assessed as Least Concern with an overall stable current population trend, but is undergoing declines in parts of its range due to hunting (Anderwald et al. 2020).

In general, our results reiterate the broad extent of use of wild species. Across the assessed groups, a predominant form of use is for pets, display and / or horticultural use, followed by hunting or collection for food. Among birds, the primary factor explaining the predominance of pets is the live cage-bird trade, which has emerged as a major driver of declines among passerines, particularly in Southeast Asia (e.g., Eaton et al. 2017). Meanwhile, cacti have long been sought after for the horticultural trade and by private collectors for their ornamental value and their perceived rarity, with both seeds and mature individuals collected (Goettsch et al. 2015). Since use and trade is not consistently recorded on the Red List, and especially so for non-threatened species, we cannot be conclusive about the full extent of use or the prevalence of different types of use in all comprehensively assessed taxonomic groups. However, our initial results confirm some taxon-specific investigations into the use of wild species, such as trees which are often used for timber as well as horticultural purposes (e.g., Beech et al. 2017).

While our results suggest that more species are *not* being impacted detrimentally by use than are, we also show that intentional biological resource use is a major threat, contributing towards increased risks of extinction for more than a quarter of Near Threatened and threatened species. The proportion of species negatively impacted by biological resource use is generally higher in aquatic taxa than among terrestrial taxa. While the impact of fisheries is well established for bony and cartilaginous fishes (Dulvy et al. 2014; MacNeil et al. 2020), the high proportion of corals and cone snails adversely impacted by biological resource use can generally be explained by increasing removals and harvests of corals for display in aquariums and for the curio-trade in the former (Bruckner 2000, Cannas et al. 2019), and by bioprospecting for conotoxin research and shell collecting in the latter (Peters et al. 2013).

Perhaps the starkest result of our analyses is that many species that are impacted by biological resource use are not currently documented as receiving any management actions that directly address this threat. The relatively high proportion of species subject to international trade controls can be explained by the fact that the most common management action that is documented is listing in a CITES Appendix. All cycads, for example, are included on CITES Appendix II through a higher-taxon listing (representing 229 out of the 255 threatened or NT cycad species in this analysis). Very few species have a national harvest management plan in place, although these appear to be more readily available for aquatic species, such as cartilaginous fishes, which have traditionally been under-represented in CITES. The high numbers of species negatively affected by use stresses the value of national management plans being in place. Of course, many species impacted by biological resource use may benefit from conservation actions that we did not directly investigate, such as the establishment and management of protected areas, community-based resource management, and other site-based interventions, while some are subject to measures to reduce demand.

There are several important caveats to our analyses. First, our study focuses on direct and intentional forms of biological resource use, but the impacts of use extend well beyond the impacts on the target species. The most evident examples of this are deforestation (specifically logging) in the terrestrial realm and by-catch in the aquatic realm. While logging is clearly a major direct threat to timber species, it can also have severe repercussions on forest-dependent species. For example, some 55% of NT and threatened bird species are affected by the unintentional effects of logging (IUCN 2020). Likewise, while commercial fishing is a direct threat to many target fisheries, by-catch is a major recognized threatening process in the sea (Komoroske & Lewison 2015). Our analysis only included by-catch for cartilaginous fishes where parts (e.g., fins, gills) of by-caught species frequently enter trade.

Second, we focused our analysis on 13 taxonomic groups that have been comprehensively assessed on the Red List, but our estimates of the extent of use of selected reptiles and dicotyledonous plants may be inflated because the families and orders included in our analysis are not necessarily representative of the broader diversity in the class (and are possibly more likely to be used). We also excluded DD and EX/EW species throughout our analyses. For DD species, assessors were not able to assign a category of extinction risk due to uncertainty in the assessment, including on the severity of threatening processes (Bland et al. 2017), such as over-exploitation; while many DD species may prove to be threatened, some have also been shown to be more widely distributed or common than previously understood (Butchart & Bird 2010). Unsustainable exploitation is already understood to have driven many species to extinction, such as the Dodo *Raphus cucullatus* and Steller’s Sea Cow *Hydrodamalis gigas*. At least 12% (102) of species listed as recently Extinct (since 1500) on the Red List have intentional use indicated as a threat that led to the species’ extinction (IUCN 2020).

Third, our study is dependent on the information captured under the IUCN Classification Schemes. Because some information is indicated as “required” (i.e. mandatory) while some is “recommended” (i.e. not mandatory), our analyses are constrained by the degree to which this information has been recorded consistently. Even where the information has been recorded, assessors may not always be aware of the full range of threats, uses or actions that apply. For example, in completing the Use and Trade Classification Scheme, full consideration may not always be given to traditional or indigenous uses; IUCN is currently preparing guidelines that would help assessors take such uses into account. Lastly, we reiterate again that 12% of the assessments included in our analysis are more than 10 years out of date, and hence the extinction risk assessments and supporting information for them may no longer be current.

In light of our caveats, we propose a few recommendations which would help improve the utility of Red List data for future analyses (Table 2). Specifically, these recommendations focus on: i) a small change to the Threats Classification Scheme, under Class 5, to indicate where the motivation for use is known, but the scale is not; ii) making the coding of scope and severity of threats in the Threats Classification Scheme recommended documentation for all threatened or NT species; iii) making documentation of threats recommended information for LC species; iv) ensuring that all assessments prioritized in the Red List’s strategic plan comply with recommended documentation requirements (which would have the benefit of also ensuring that conservation actions in place and needed are documented for all species); and iv) the addition of a simple check-box to indicate whether or not the classification schemes for a given species assessment have been filled in at the recommended level, which would allow one to make a rapid determination of the completeness of Red List assessment supporting documentation for the purposes of analysis. We make these recommendations recognizing the time and resource constraints in the Red List process, which is driven largely by volunteer scientists. There is a well-documented tension between the desire to expand the taxonomic breadth of assessed species with the need to undertake timely reassessments, all the while ensuring the supporting information is as complete as it can be (Rondinini et al. 2013). Given this, we cannot, for example, propose that a suite of information fields be made mandatory for assessors, while simultaneously acknowledging that many species urgently need reassessment. Further discussion on our recommendations is provided in the Supporting Information.

**Table 2.**
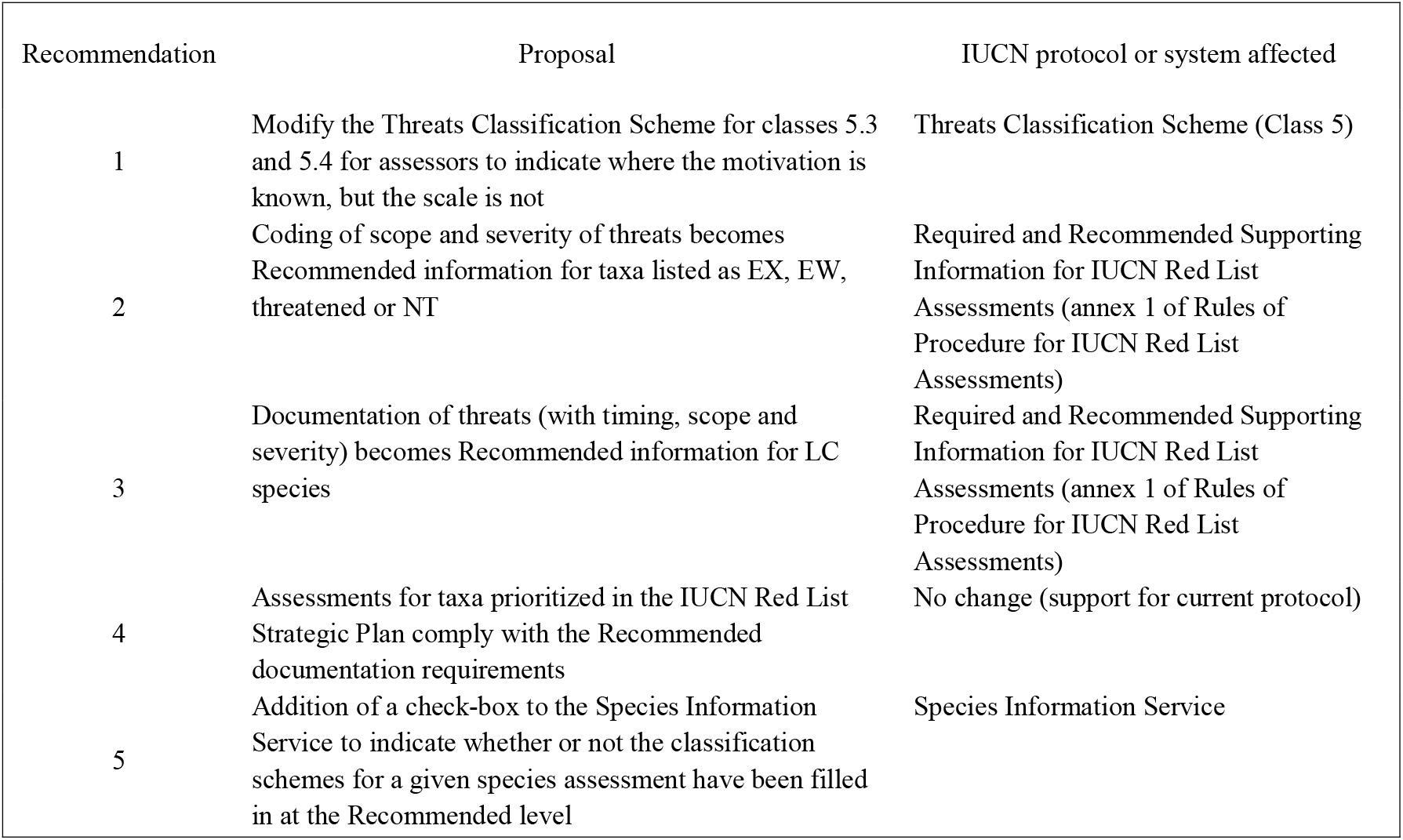
Recommendations for improving consistency and available information in use-related Red List data.

As previous studies have shown, Red List data can play a key role in supporting major global assessment processes, and by extension broader international policy. Our study uses Red List data to quantify the degree to which the use of wild species either does or does not impact negatively on species extinction risk, and thus whether documented use may be biologically sustainable or unsustainable. Our ability to disentangle the nature and extent of this use could be considerably improved through some minor amendments to established Red List protocols and through greater efforts by assessors and Red List assessment initiatives to ensure, wherever possible, more consistent recording of the information and data underpinning Red List assessments. Nonetheless, our findings show that while over-exploitation is clearly a direct major threat to many species, and indeed has already driven some to extinction, there are also many species for which use is currently taking place at levels that are unlikely to contribute significantly to an increase in their extinction risk.

Our results can usefully inform processes such as the IPBES assessment on sustainable use, and our analysis provides important information for international policy-making, since the Red List is frequently taken into consideration in policy deliberations such as the progress of species through the CITES Review of Significant Trade (RST) process (CITES 2018). However, there is no reason why these data could not also be harvested to inform other decision-making, both within but also beyond CITES. Our study provides a first approach for disentangling likely sustainable versus likely unsustainable use of species from the Red List; ideally this information should be combined with other data sources, including from national or regional datasets, to provide the most complete picture on the impacts of use of wild species as possible. Clearly, more effort needs to be invested in understanding the factors that determine whether use is sustainable, and the effectiveness of different mitigation actions.

## Acknowledgments

We thank Kathryn Philipps for helpful inputs into the manuscript. This manuscript was prepared with funding support from the French Ministry for Ecological Transition (MTE). IUCN thanks the MTE for globally supporting IUCN’s engagement with IPBES in the frame of the IUCN-France Partnership. MB is supported by a generous grant from the Rufford Foundation. DWSC acknowledges funding from the UK Research and Innovation’s Global Challenges Research Fund (UKRI GCRF) through the Trade, Development and the Environment Hub project (project number ES/S008160/1). The views expressed in this publication do not necessarily reflect those of IUCN. The designation of geographical entities in this paper, and the presentation of the material, do not imply the expression of any opinion whatsoever on the part of IUCN concerning the legal status of any country, territory, or area, or of its authorities, or concerning the delimitation of its frontiers or boundaries.

## Notes

### Competing Interest Statement

The authors have declared no competing interest.

### Summary of Updates

This manuscript has been revised to clarify methods and update Supplemental files.

