## Supporting Information for "Prevalence of sustainable and unsustainable use of wild species inferred from the IUCN Red List"

*Glossary of Red List-related terms used in the manuscript*

*Biological resource use*: threats from consumptive use of “wild” biological resources including deliberate and unintentional harvesting effects; also persecution or control of specific species (Salafsky et al. 2008). Part of the IUCN-Conservation Measures Partnership (CMP) classification of direct threats to biodiversity and the IUCN Red List Threat Classification Scheme (Class 5). See Table S2.

*Comprehensively assessed / comprehensive assessment:* taxonomic groups that include at least 150 species, of which >80% have been assessed (IUCN 2020).

*Conservation Actions In Place Classification Scheme:* Used by assessors to record what conservation actions are already in place for a taxon. This information is not mandatory, only recommended. For definitions, examples and guidance on the Conservation Actions In Place Classification Scheme v. 2.0, see <https://nc.iucnredlist.org/redlist/content/attachment_files/dec_2012_guidance_conservation_actions_in_place_classification_scheme.pdf> .

*Current population trend*: Current population trend refers to trends over a period of ca. three years around the date the assessment was done; can be set by assessors as “Increasing”, “Decreasing”, “Stable”, or “Unknown”.

*Data sufficient species:* Excludes species listed as Data Deficient on the IUCN Red List.

*Harvest management plan:* a plan exists for the species, or populations of the species, which sets out levels of (sustainable) harvest; assessors can select “Yes”, “No” or “Unknown”. This information is not mandatory, only recommended.

*International management or trade controls*: Examples include CITES, Regional Fisheries Agreements, ICCAT, Marine Stewardship Council, Forestry Stewardship Council, Marine Aquarium Council, Phytosanitary Measures Agreement, US Endangered Species Act, etc. (see <https://nc.iucnredlist.org/redlist/content/attachment_files/dec_2012_guidance_conservation_actions_in_place_classification_scheme.pdf>). Assessors can select “Yes”, “No” or “Unknown”. This information is not mandatory, only recommended.

*Recommended documentation*: Recommended supporting information is not essential for a Red List assessment to be accepted for publication on the IUCN Red List but is strongly encouraged for all assessments for taxa prioritized in the IUCN Red List Strategic Plan. See: <https://nc.iucnredlist.org/redlist/content/attachment_files/Required_and_Recommended_Supporting_Information_for_IUCN_Red_List_Assessments.pdf>.

*Mandatory documentation*: Supporting information that is required (i.e., mandatory) for ALL Red List assessments before they can be accepted for publication on the IUCN Red List. See: <https://nc.iucnredlist.org/redlist/content/attachment_files/Required_and_Recommended_Supporting_Information_for_IUCN_Red_List_Assessments.pdf>.

*Threatened:* Taxa listed as Vulnerable (VU), Endangered (EN), or Critically Endangered (CR) on the IUCN Red List.

*Threats Classification Scheme*: Used by assessors to record past, ongoing and future threats to a taxon. Documenting major threats impacting a species is mandatory for species listed as EX, EW, threatened, and NT. For definitions, examples and guidance on the Threats Classification Scheme ver. 3.2, see: <https://nc.iucnredlist.org/redlist/content/attachment_files/dec_2012_guidance_threats_classification_scheme.pdf>

*Threat impact score*: A score calculated from a combination of threat scope, severity and timing. See Table S5.

*Threat scope*: The scope of the threat, given as: the threat affects the whole (>90%); majority (50-90%); minority (<50%); or an unknown proportion of the population. This is only optional information to be provided in the Red List assessment.

*Threat severity*: The severity of the threat, given as: causing very rapid declines; rapid declines; slow, significant declines; causing/could cause fluctuations; negligible declines; no decline; or unknown. This is only optional information to be provided in the Red List assessment.

*Threat timing*: The timing of the threat given as: ongoing; past (unlikely to return); past (likely to return); future; or unknown. This is recommended information to be provided in the Red List assessment.

*Use and Trade Classification Scheme*: Used by assessors to record how a taxon is utilized and what level of trade occurs for the taxon. This information is not mandatory, only recommended. For the General Use and Trade Classification Scheme (including the Non-Consumptive Use scheme) ver. 1.0, see <https://nc.iucnredlist.org/redlist/content/attachment_files/July_2020_Guidance_General_Use_and_Trade_Classification_Scheme.pdf>. See Table S3.

Supplementary Methods

*The main purposes of use of wild animal and plant species recorded in the Red List*

We investigated the prevalence of different purposes of use from the use and trade information recorded in the Red List. Because completing the Use and Trade classification scheme is not mandatory for Red List assessors, we investigated the prevalence of use and trade recording to decide which species groups to include in our analyses. We selected taxonomic groups for inclusion based on the following criteria: i) >40% of all extant, data sufficient species have at least one purpose of use coded (thus selecting taxonomic groups with high prevalence of use); and / or ii) the proportion of LC species with at least one purpose of use code falls above or within the range of the proportion of species with use and trade recording across the other Red List categories (thus also selecting taxonomic groups where use and trade may be relatively low, but use and trade in LC species is coded to a similar level as that of species in other RL categories). This limited our dataset to the following taxonomic groups which have adequate recording of use and trade: birds, amphibians, selected reptiles, cycads, conifers and selected dicots from the terrestrial group; and corals, selected bony fishes, crustaceans and cone snails from the aquatic species group (Table S4). We excluded mammals, cephalopods and cartilaginous fishes as they are meeting neither criteria i) nor ii), meaning at the level of each of these three groups there is either relatively little use or documentation of use in the Red List is incomplete.

For each taxonomic group, we calculated the total number of species recorded as being used for at least one purpose in the Use and Trade classification scheme. However, we excluded those cases where species were used for establishing ex-situ production (use code 16), “other” (17), and where purpose of use was unknown (18). Ex-situ production involves establishing captive populations for conservation breeding and translocation but also for ranching, farming or propagation purposes; unfortunately, it is not possible to distinguish these purposes without assessors voluntarily providing information on harvest from captive and cultivated sources. Only 11 species (six cartilaginous fishes, four bony fishes, and one bird) in our analyses had an “unknown” use and trade category. We summarized the data as the percentage of species recorded for different types of use on the Red List.

*Wild species for which intentional use is having a negative impact*

Since not all types of biological resource use are directly targeted at the species in question, and hence immediately relevant to our analyses, we developed a decision-tree (Figure S1) for removing those types of threats that are not relevant to an analysis of sustainable and unsustainable direct use of species.

First, we discounted the threat of persecution or control, as this generally does not represent intentional consumptive use of wild species. We also excluded those threats where the species was not the deliberate target of the activity, here termed unintentional use, with the exception of those groups of aquatic species which are widely used even if caught unintentionally as bycatch (e.g., cartilaginous fishes; Dulvy et al. 2014). Any records of vertebrates documented as threatened by intentional gathering of plants or logging and wood harvesting were also excluded as these likely represent coding errors by the assessors and should have been coded as unintentional (i.e., the species is not the direct target of the activity).

In addition to intentional and unintentional use, the IUCN Threat Classification scheme also allows for biological resource use to be recorded if the motivation behind it is unknown (i.e., it is unknown if use is intentionally targeting the species in question or if it is threatening it unintentionally). While intentional gathering of terrestrial plants or logging and wood harvesting (threat codes 5.2.4 and 5.3.5) can threaten animals through habitat loss, they do not directly lead to the use of animals and we excluded any such records for animal groups. Where biological resource use with unknown motivation was recorded for species (codes 5.1.4 and 5.4.6 for animal species; codes 5.2.4 and 5.3.5 for plant species; Table S2), we reviewed the text of individual species assessments to determine whether these records should be excluded from the analysis. For instance, the Red List documentation states that the Pygmy Slow Loris (*Nycticebus pygmaeus*) is recorded as threatened by intentional hunting and collecting of terrestrial animals (5.1.1), and by fishing and harvesting of aquatic resources for which the motivation is unknown or unrecorded (5.4.6). This terrestrial species is exploited for the pet trade and harvested for medicinal use, hunted as a food source, and experiences habitat loss due to agriculture. Its categorization under 5.4.6 is likely to represent a threat coding error and was removed, as no aquatic resource use is known to directly affect this terrestrial species. For other biological use threats with unknown motivation, we were unable to determine whether the threat was directly targeting the species or not. We present this uncertainty in our results as a range where the minimum proportion includes all species with threats that could be conclusively determined as intentional (and hence is more evidentiary), and the maximum proportion additionally includes those species with motivation unknown or unrecorded that may represent further cases of intentional biological resource use (and hence is more precautionary). For groups where no species have such motivation unknown / unrecorded threats, we present only the minimum.

We only included biological resource use if it had a major impact on species survival. The Red List uses a scoring system to derive threat impact, based on three key elements: timing of the threat, categorized as past (unlikely to return), past (likely to return), ongoing, future, or unknown; scope of the threat, categorized as the percentage of the population affected by the threat, i.e., whole, >90%, majority, 50-90%, minority, <50%, or unknown; and threat severity, categorized as causing very rapid declines, rapid declines, slow and significant declines, causing/would cause fluctuations or negligible declines, or unknown. This information is used by the Red List to create an overall threat impact score by summing the scores for timing, scope and severity for each threat impacting a species (IUCN 2020). While threat timing is recommended information to be provided in the Red List assessment (i.e., strongly encouraged but not essential to publication), severity and scope are entirely optional. We thus utilized this impact score categorization, but with several amendments (Table S5). Threats classed as in the past but likely to return were treated the same as future threats, receiving a score of 1; threats with either unknown or missing timing, severity and scope information were assigned a medium score of 2. Subsequently, threats with a threat impact score showing low, negligible or no impact were excluded, and medium to high impact threats (threat impact score >= 6) were retained in our analysis. As an example, the amphibian *Herpele squalostoma* is threatened by intentional use (5.4.2, where the species is the target): this is an ongoing threat (timing) associated with a minority scope (<50% of population affected), causing negligible declines (severity). We calculated an impact score of 4 and thus excluded this particular threat from analysis. The Northern Fur Seal *Callorhinus ursinus* was threatened by large-scale intentional use (5.4.2) in the past (it is unlikely to return) a threat of majority scope, possibly causing fluctuations. This threat also yielded an impact score of 4 and was excluded as well. The main implication of our amended scoring is that it is precautionary in including threats where both severity and scope are marked as unknown (as medium impact), but evidentiary in excluding threats if at least one of them is coded as either negligible or minority.

*Summary of analyses undertaken*

We provide a short-hand summary of the main analyses undertaken in this paper, which correspond to the numbers at the bottom of Table 1.

1. The main purposes of use of wild animal and plant species on the Red List

Estimated as: the total number of species recorded as being used for at least one purpose of use as the percentage of extant, data sufficient species recorded for different types of use (Fig. 1)

2. Wild species for which intentional use is having a negative impact on extinction risk (i.e., for which “biological resource use” is documented as a major threat), estimated from among:

a. all extant, data-sufficient species with at least one purpose of use coded (sensu analysis 1);
b. all NT and threatened species (from among all 13 taxonomic groups comprehensively assessed; Fig. 2, S2)

3. Wild species for which intentional use is not having a negative impact on extinction risk. Estimated from the number of extant, data sufficient species recorded as being subject to some form of use or trade that:

a. are currently LC;
b. currently LC and not declining (i.e., have either stable or increasing current population trends; Fig. 3);
c. are threatened or NT and are not documented as having intentional use as a major threat and have stable or increasing population trends.

4. Conservation actions in place or lacking for utilized wild species.

Estimated from the number of NT and threatened species that are adversely affected by biological resource use and:

a. are coded as receiving either one or both of international trade controls and / or harvest management actions (Fig. 4);
b. are not receiving either of these conservation management actions.

Supplementary Discussion

*Recommendations for improving consistency and available information in use-related Red List data*

In this paper, we propose a few recommendations (Table 2) that would help to reduce the proportion of used species – currently nearly half – for which we have no evidence as to whether intentional use is sustainable or not. As discussed in the main text, we do not make these recommendations lightly, given the delicate balance between expanding the taxonomic breadth of the Red List with the best possible supporting information and the need to undertake timely reassessments and the demand on the time and resources of individual assessors. We do however consider our recommendations achievable.

Recommendation one concerns coding of the threat category “motivation unknown”. In the current Threats Classification Scheme, if the assessor does not know the scale of harvest (i.e., whether “subsistence/small scale” or “large scale”), their only option is to select the option for “motivation unknown / unrecorded” under the sections 5.3 (logging & wood harvesting) and 5.4. (fishing & harvesting aquatic resources), even though they are likely to know whether the use is intentional or unintentional. For instance, we found 1,519 amphibian species for which “logging and wood harvesting” was coded as a threat, of which 1,187 species threats were recorded under “motivation unknown”, but the motivation must have been unintentional as these species are impacted through the loss of forest, not through direct exploitation. Although our methodology excludes these cases from our analyses, we propose a modification to the Threats Classification Scheme for assessors to indicate where the motivation is known, but the scale is not, to avoid these coding issues in the future (Table S2).

Recommendation two is that data on timing, scope and severity of threats should be better coded as it would allow us to tease out more effectively where threat impacts are medium to high and bring greater precision to our results. For corals and cone snails, the threat of biological resource use is likely to be small in comparison with the impacts of bleaching and disease for corals (Carpenter et al. 2008), and urban pollution, tourism and coastal development for cone snails (Peters et al. 2013).

Further, it would be useful to better understand and quantify the degree to which species can be subject to some level of use without this resulting in them becoming threatened (i.e., impact is low, highly localized, negligible or no impact). This requires that the effects of biological resource use be more consistently recorded for LC species (Recommendation three).

Recommendation four is for all assessments in the Red List Strategic Plan to comply with the recommended documentation requirements. One further advantage this would offer is that it would mean better availability of information on conservation actions in place and needed, respectively. Indeed, our results based on analysis of data in the Conservation Actions in Place Classification Scheme are particularly constrained because data are available for a limited number of species. For example, only one taxonomic group (conifers) has more than half of its NT and threatened species with some documentation of whether a harvest management plan is in place (as opposed to being left blank). Whether international trade controls are in place is generally better documented than whether a harvest management plan is in place. This is likely because information on whether species are in a CITES Appendix or subject to some other policy controls is easier to obtain than whether a harvest management plan is in place.

Finally, the addition of a check-box to indicate when the classification schemes for a given species assessment have been filled in at the recommended level, would be a powerful addition to the Red List documentation (Recommendation five).

Supplementary Tables

Table S1. Taxonomic description of species in comprehensively assessed groups of the Red List.

| **Taxonomic group** | **Phylum** | **Class** | **Order** | **Family** | **Genus** |
| --- | --- | --- | --- | --- | --- |
| Cephalopods | Mollusca | Cephalopoda* | All species | | |
| Cone snails | Mollusca | Gastropoda | Neogastropoda | Conidae | Conus |
| Corals | Cnidaria | Hydrozoa | All species | | |
|  |  | Anthozoa | Helioporacea | All species | |
|  |  |  | Scleractinia | Acroporidae | All species |
|  |  |  |  | Agariciidae | All species |
|  |  |  |  | Astrocoeniidae | All species |
|  |  |  |  | Euphyllidae | All species |
|  |  |  |  | Faviidae | All species |
|  |  |  |  | Mussidae | All species |
|  |  |  |  | Oculinidae | All species |
|  |  |  |  | Pectiniidae | All species |
|  |  |  |  | Pocilloporidae | All species |
|  |  |  |  | Poritidae | All species |
|  |  |  |  | Rhizangiidae | All species |
|  |  |  |  | Siderastreidae | All species |
|  |  |  |  | Trachyphylliidae | All species |
|  |  |  |  | Turbinoliidae | All species |
|  |  |  |  | Dendrophylliidae | *Balanophyllia* |
|  |  |  |  |  | *Duncanopsammia* |
|  |  |  |  |  | *Heteropsammia* |
|  |  |  |  |  | *Turbinaria* |
|  |  |  |  | Caryophylliidae | *Heterocyathus* |
| Cartilaginous fishes (sharks, rays, chimaeras) | Chordata | Chondrichthyes | All species | | |
| Bony fishes (selected): |  |  |  |  |  |
| Sturgeons | Chordata | Actinopterygii | Acipenseriformes | All species | |
| Tarpons & bonefishes |  |  | Albuliformes | All species | |
|  |  |  | Elopiformes | All species | |
| Anchovies, sardines etc. |  |  | Clupeiformes | All species | |
| Groupers & wrasses |  |  | Perciformes | Epinephelidae | All species |
|  |  |  |  | Labridae | All species |
| Tunas |  |  |  | Scombridae | All species |
| Billfishes |  |  |  | Istiophoridae | All species |
|  |  |  |  | Xiphiidae | All species |
| Blennies |  |  |  | Blenniidae | All species |
|  |  |  |  | Chaenopsidae | All species |
|  |  |  |  | Clinidae | All species |
|  |  |  |  | Dactyloscopidae | All species |
|  |  |  |  | Labrisomidae | All species |
|  |  |  |  | Tripterygiidae | All species |
| Seabreams |  |  |  | Sparidae | All species |
|  |  |  |  | Centracanthidae | All species |
| Angelfishes |  |  |  | Pomacanthidae | All species |
| Butterflyfishes |  |  |  | Chaetodontidae | All species |
| Surgeonfishes |  |  |  | Acanthuridae | All species |
| Pufferfishes |  |  | Tetraodontiformes | Tetraodontidae | All species |
| Seahorses & pipefish |  |  | Syngnathiformes | Syngnathidae | All species |
| Trumpetfishes |  |  |  | Aulostomidae | All species |
| Shrimpfishes |  |  |  | Centriscidae | All species |
| Seamoths |  |  |  | Pegasidae | All species |
| Ghost pipefishes |  |  |  | Solenostomidae | All species |
| Cornetfishes |  |  | Gasterosteiformes | Fistulariidae | All species |
| Crustaceans (selected): |  |  |  |  |  |
| Lobsters | Arthropoda | Malacostraca | Decapoda | Glypheidae | All species |
|  |  |  |  | Polychelidae | All species |
|  |  |  |  | Nephropidae | All species |
|  |  |  |  | Enoplometopidae | All species |
|  |  |  |  | Palinuridae | All species |
|  |  |  |  | Scyllaridae | All species |
| Freshwater crayfishes |  |  |  | Astacidae | All species |
|  |  |  |  | Cambaridae | All species |
|  |  |  |  | Parastacidae | All species |
| Freshwater crabs |  |  |  | Trichodactylidae | All species |
|  |  |  |  | Potamidae | All species |
|  |  |  |  | Potamonautidae | All species |
|  |  |  |  | Gecarcinucidae | All species |
|  |  |  |  | Pseudothelphusidae | All species |
| Freshwater shrimps |  |  |  | Alpheidae | FW species only |
|  |  |  |  | Atyidae | FW species only |
|  |  |  |  | Desmocarididae | FW species only |
|  |  |  |  | Euryrhynchidae | FW species only |
|  |  |  |  | Palaemonidae | FW species only |
|  |  |  |  | Typhlocarididae | FW species only |
|  |  |  |  | Xiphocarididae | FW species only |
| Amphibians | Chordata | Amphibia | All species | | |
| Birds | Chordata | Aves | All species | | |
| Mammals | Chordata | Mammalia | All species | | |
| Conifers | Tracheophyta | Pinopsida | All species | | |
| Cycads |  | Cycadopsida | All species | | |
| Dicotyledons (selected): |  |  |  |  |  |
| Cacti | Tracheophyta | Magnoliopsida | Caryophyllales | Cactaceae | All species |
| Magnolias |  |  | Magnoliales | Magnoliaceae | All species |
| Birches |  |  | Fagales | Betulaceae | All species |
| Southern beeches |  |  |  | Nothofagaceae | All species |
| Teas |  |  | Ericales | Theaceae | All species |
| Reptiles (selected): |  |  |  |  |  |
| Chameleons | Chordata | Reptilia | Squamata | Chamaeleonidae | All species |
| Sea snakes |  |  |  | Homalopsidae | All species |
|  |  |  |  | Elapidae | *Aipysurus* |
|  |  |  |  |  | *Emydocephalus* |
|  |  |  |  |  | *Ephalophis* |
|  |  |  |  |  | *Hydrelaps* |
|  |  |  |  |  | *Hydrophis* |
|  |  |  |  |  | *Kerilia* |
|  |  |  |  |  | *Kolpophis* |
|  |  |  |  |  | *Laticauda* |
|  |  |  |  |  | *Parahydrophis* |
|  |  |  |  |  | *Thalassophis* |
|  |  |  |  | Acrochordidae | *Acrochordus* |
|  |  |  |  | Natricidae | *Anoplohydrus* |
| Crocodiles & alligators |  |  | Crocodylia | All species | |
| Marine turtles |  |  | Testudines | Cheloniidae | All species |
|  |  |  |  | Dermochelyidae | All species |

*Nautiluses are not yet assessed

Table S2. A) The IUCN Red List Threats Classification Scheme Class 5 following Salafsky et al. (2008), with B) Proposed amendments to the scheme to account for instances where scale of use is unknown versus motivation. Bold denotes changes.

A)

| **Threats 5. Biological Resource Use** | |
| --- | --- |
| 5.1 | Hunting & collecting terrestrial animals  5.1.1 Intentional use (species being assessed is the target)  5.1.2 Unintentional effects (species being assessed is not the target)  5.1.3 Persecution/control  5.1.4 Motivation unknown/unrecorded |
| 5.2 | Gathering terrestrial plants  5.2.1 Intentional use (species being assessed is the target)  5.2.2 Unintentional effects (species being assessed is not the target)  5.2.3 Persecution/control  5.2.4 Motivation unknown/unrecorded |
| 5.3 | Logging & wood harvesting  5.3.1 Intentional use: subsistence/small scale (species being assessed is the target)  5.3.2 Intentional use: large scale (species being assessed is the target)  5.3.3 Unintentional effects: subsistence/small scale (species being assessed is not the target)  5.3.4 Unintentional effects: large scale (species being assessed is not the target)  5.3.5 Motivation unknown/unrecorded |
| 5.4 | Fishing & harvesting aquatic resources  5.4.1 Intentional use: subsistence/small scale (species being assessed is the target)  5.4.2 Intentional use: large scale (species being assessed is the target)  5.4.3 Unintentional effects: subsistence/small scale (species being assessed is not the target)  5.4.4 Unintentional effects: large scale (species being assessed is not the target)  5.4.5 Persecution/control  5.4.6 Motivation unknown/unrecorded |

B)

| **Threats 5. Biological Resource Use** | |
| --- | --- |
| 5.3 | Logging & wood harvesting  5.3.1 Intentional use: subsistence/small scale (species being assessed is the target)  5.3.2 Intentional use: large scale (species being assessed is the target)  **5.3.3 Intentional use: scale unknown (species being assessed is the target) [harvest]**  **5.3.4** Unintentional effects: subsistence/small scale (species being assessed is not the target)  **5.3.5** Unintentional effects: large scale (species being assessed is not the target)  **5.3.6 Unintentional effects: scale unknown (species being assessed is not the target) [harvest]**  **5.3.7** Motivation unknown/unrecorded |
| 5.4 | Fishing & harvesting aquatic resources  5.4.1 Intentional use: subsistence/small scale (species being assessed is the target)  5.4.2 Intentional use: large scale (species being assessed is the target)  **5.4.3 Intentional use: scale unknown (species being assessed is the target) [harvest]**  **5.4.4** Unintentional effects: subsistence/small scale (species being assessed is not the target)  **5.4.5** Unintentional effects: large scale (species being assessed is not the target)  **5.4.6 Unintentional effects: scale unknown (species being assessed is not the target) [harvest]**  **5.4.7** Persecution/control  **5.4.8** Motivation unknown/unrecorded |

Table S3. Red List Use and Trade classification scheme. Analyses in this paper exclude use codes 16 – 18.

| 1. Food – human 2. Food – animal 3. Medicine – human & veterinary 4. Poisons 5. Manufacturing chemicals 6. Other chemicals 7. Fuels 8. Fibre 9. Construction or structural materials | 10. Wearing apparel, accessories 11. Other household goods 12. Handicrafts, jewellery, etc. 13. Pets / display animals, horticulture 14. Research 15. Sport hunting / specimen collecting 16. Establishing ex situ production 17. Other (free text) 18. Unknown |
| --- | --- |

Table S4. Proportion of extant, data sufficient species in each taxonomic group with at least one purpose of use documented on the Red List. ^[1]^

| **Taxonomic group** | **Proportion LC** | **Proportion threatened** | **Proportion all species** | **LC in range** | **Criterion i** | **Criterion ii** |
| --- | --- | --- | --- | --- | --- | --- |
| Amphibians | 0.12 | 0.09 | 0.11 | Within | No | Yes |
| Birds | 0.46 | 0.47 | 0.46 | Within | Yes | Yes |
| Selected bony fishes | 0.53 | 0.48 | 0.53 | Within | Yes | Yes |
| Cartilaginous fishes | 0.22 | 0.51 | 0.35 | Below | No | No |
| Cephalopods | 0.20 | 0.20 | 0.21 | Below | No | No |
| Cone snails | 1.00 | 0.98 | 1.00 | Within | Yes | Yes |
| Conifers | 0.82 | 0.69 | 0.76 | Above | Yes | Yes |
| Corals | 0.67 | 0.79 | 0.75 | Within | Yes | Yes |
| Crustaceans | 0.17 | 0.11 | 0.15 | Above | No | Yes |
| Cycads | 0.69 | 0.85 | 0.80 | Below | Yes | No |
| Selected dicots | 0.59 | 0.58 | 0.58 | Within | Yes | Yes |
| Mammals | 0.21 | 0.43 | 0.28 | Below | No | No |
| Selected reptiles | 0.53 | 0.41 | 0.46 | Above | Yes | Yes |

Table S5. Scoring system for the impact of threats used in the current analysis, adapted from the Red List.^[2]^

| Timing:  Ongoing threat (+3) | | | | | Future / Past, Likely to Return threat (long term) (+1) | | | |
| --- | --- | --- | --- | --- | --- | --- | --- | --- |
| Severity:  Scope: | *Very rapid*  (+3) | *Rapid / Unknown*  (+2) | *Slow / Fluctuating*  (+1) | *Negligible / No impact* (0) | *Very rapid*  (+3) | *Rapid / Unknown*  (+2) | *Slow / Fluctuating*  (+1) | *Negligible / No impact* (0) |
| Whole (+3) | 9 | 8 | 7 | 6 | 7 | 6 | 5 | 4 |
| Majority / Unknown (+2) | 8 | 7 | 6 | 5 | 6 | 5 | 4 | 3 |
| Minority (+1) | 7 | 6 | 5 | 4 | 5 | 4 | 3 | 2 |

Additive impact scores:

8-9: High impact threat

6-7: Medium impact threat

3-5: Low impact threat

0-2: Negligible / no impact threat

Table S7. Threatened or NT species with at least one purpose of use or trade recorded that are not impacted by major, intentional biological resource use, and have stable or increasing population trends.

| **Taxonomic group** | **Scientific name** | **Red List Category** | **Population trend** |
| --- | --- | --- | --- |
| Amphibians | *Leptobrachium ailaonicum* | Near Threatened | Stable |
| Amphibians | *Atelopus flavescens* | Vulnerable | Stable |
| Amphibians | *Eurycea rathbuni* | Vulnerable | Stable |
| Amphibians | *Lyciasalamandra helverseni* | Vulnerable | Stable |
| Amphibians | *Lyciasalamandra luschani* | Vulnerable | Stable |
| Amphibians | *Lyciasalamandra atifi* | Endangered | Stable |
| Birds | *Accipiter collaris* | Near Threatened | Stable |
| Birds | *Accipiter poliogaster* | Near Threatened | Increasing |
| Birds | *Acrocephalus rodericanus* | Near Threatened | Increasing |
| Birds | *Actinodura sodangorum* | Near Threatened | Stable |
| Birds | *Anas chlorotis* | Near Threatened | Increasing |
| Birds | *Buteo solitarius* | Near Threatened | Stable |
| Birds | *Callaeas wilsoni* | Near Threatened | Increasing |
| Birds | *Caprimulgus phalaena* | Near Threatened | Stable |
| Birds | *Charadrius aquilonius* | Near Threatened | Increasing |
| Birds | *Charadrius melodus* | Near Threatened | Increasing |
| Birds | *Charadrius pallidus* | Near Threatened | Stable |
| Birds | *Chloebia gouldiae* | Near Threatened | Stable |
| Birds | *Cynanthus lawrencei* | Near Threatened | Stable |
| Birds | *Dendrocopos owstoni* | Near Threatened | Stable |
| Birds | *Dendropicos stierlingi* | Near Threatened | Stable |
| Birds | *Ducula whartoni* | Near Threatened | Stable |
| Birds | *Egretta rufescens* | Near Threatened | Increasing |
| Birds | *Elanus scriptus* | Near Threatened | Stable |
| Birds | *Erythrura coloria* | Near Threatened | Stable |
| Birds | *Estrilda poliopareia* | Near Threatened | Stable |
| Birds | *Eudyptes schlegeli* | Near Threatened | Stable |
| Birds | *Euplectes jacksoni* | Near Threatened | Stable |
| Birds | *Foudia flavicans* | Near Threatened | Increasing |
| Birds | *Fringilla teydea* | Near Threatened | Increasing |
| Birds | *Garrulax nuchalis* | Near Threatened | Stable |
| Birds | *Gyps himalayensis* | Near Threatened | Stable |
| Birds | *Hemiphaga novaeseelandiae* | Near Threatened | Increasing |
| Birds | *Hypsipetes borbonicus* | Near Threatened | Increasing |
| Birds | *Larus atlanticus* | Near Threatened | Stable |
| Birds | *Larvivora komadori* | Near Threatened | Stable |
| Birds | *Megascops marshalli* | Near Threatened | Stable |
| Birds | *Microcarbo coronatus* | Near Threatened | Stable |
| Birds | *Oxyura australis* | Near Threatened | Stable |
| Birds | *Parotia wahnesi* | Near Threatened | Stable |
| Birds | *Phalcoboenus australis* | Near Threatened | Stable |
| Birds | *Philesturnus carunculatus* | Near Threatened | Increasing |
| Birds | *Philesturnus rufusater* | Near Threatened | Increasing |
| Birds | *Phoebastria immutabilis* | Near Threatened | Stable |
| Birds | *Phoebastria nigripes* | Near Threatened | Increasing |
| Birds | *Polytelis alexandrae* | Near Threatened | Stable |
| Birds | *Psittinus abbotti* | Near Threatened | Stable |
| Birds | *Pyrilia caica* | Near Threatened | Stable |
| Birds | *Pyrrhura devillei* | Near Threatened | Stable |
| Birds | *Speculanas specularis* | Near Threatened | Stable |
| Birds | *Streptopelia reichenowi* | Near Threatened | Stable |
| Birds | *Thalasseus elegans* | Near Threatened | Stable |
| Birds | *Todiramphus pelewensis* | Near Threatened | Stable |
| Birds | *Amazona arausiaca* | Vulnerable | Increasing |
| Birds | *Amazona guildingii* | Vulnerable | Increasing |
| Birds | *Anas albogularis* | Vulnerable | Stable |
| Birds | *Anas aucklandica* | Vulnerable | Stable |
| Birds | *Anthropoides paradiseus* | Vulnerable | Stable |
| Birds | *Ardenna bulleri* | Vulnerable | Stable |
| Birds | *Branta sandvicensis* | Vulnerable | Increasing |
| Birds | *Buteo galapagoensis* | Vulnerable | Stable |
| Birds | *Capito wallacei* | Vulnerable | Stable |
| Birds | *Charadrius sanctaehelenae* | Vulnerable | Increasing |
| Birds | *Coracopsis barklyi* | Vulnerable | Stable |
| Birds | *Cyanoramphus unicolor* | Vulnerable | Stable |
| Birds | *Diomedea epomophora* | Vulnerable | Stable |
| Birds | *Eunymphicus cornutus* | Vulnerable | Increasing |
| Birds | *Eunymphicus uvaeensis* | Vulnerable | Increasing |
| Birds | *Falco araeus* | Vulnerable | Stable |
| Birds | *Falco hypoleucos* | Vulnerable | Stable |
| Birds | *Fregata aquila* | Vulnerable | Stable |
| Birds | *Fulica alai* | Vulnerable | Stable |
| Birds | *Grus monacha* | Vulnerable | Increasing |
| Birds | *Hemiphaga chathamensis* | Vulnerable | Increasing |
| Birds | *Icterus oberi* | Vulnerable | Stable |
| Birds | *Leucocarbo campbelli* | Vulnerable | Stable |
| Birds | *Leucocarbo colensoi* | Vulnerable | Stable |
| Birds | *Leucocarbo ranfurlyi* | Vulnerable | Stable |
| Birds | *Lonchura vana* | Vulnerable | Stable |
| Birds | *Megapodius pritchardii* | Vulnerable | Increasing |
| Birds | *Nannopterum harrisi* | Vulnerable | Stable |
| Birds | *Nesoenas mayeri* | Vulnerable | Stable |
| Birds | *Ninox natalis* | Vulnerable | Stable |
| Birds | *Odontophorus dialeucos* | Vulnerable | Stable |
| Birds | *Phoebastria albatrus* | Vulnerable | Increasing |
| Birds | *Phoenicoparrus andinus* | Vulnerable | Stable |
| Birds | *Pica nutalli* | Vulnerable | Stable |
| Birds | *Poicephalus robustus* | Vulnerable | Stable |
| Birds | *Procellaria parkinsoni* | Vulnerable | Stable |
| Birds | *Psittacula eques* | Vulnerable | Increasing |
| Birds | *Pterodroma axillaris* | Vulnerable | Increasing |
| Birds | *Pterodroma deserta* | Vulnerable | Stable |
| Birds | *Pterodroma solandri* | Vulnerable | Increasing |
| Birds | *Pyrrhula murina* | Vulnerable | Stable |
| Birds | *Pyrrhura perlata* | Vulnerable | Stable |
| Birds | *Scolopax mira* | Vulnerable | Stable |
| Birds | *Thalassarche eremita* | Vulnerable | Stable |
| Birds | *Touit huetii* | Vulnerable | Stable |
| Birds | *Alopecoenas rubescens* | Endangered | Stable |
| Birds | *Copsychus sechellarum* | Endangered | Increasing |
| Birds | *Cyclopsitta coxeni* | Endangered | Stable |
| Birds | *Foudia aldabrana* | Endangered | Increasing |
| Birds | *Fringilla polatzeki* | Endangered | Increasing |
| Birds | *Grus americana* | Endangered | Increasing |
| Birds | *Hypotaenidia sylvestris* | Endangered | Stable |
| Birds | *Junco insularis* | Endangered | Increasing |
| Birds | *Nipponia nippon* | Endangered | Increasing |
| Birds | *Ognorhynchus icterotis* | Endangered | Increasing |
| Birds | *Papasula abbotti* | Endangered | Stable |
| Birds | *Platalea minor* | Endangered | Increasing |
| Birds | *Porphyrio hochstetteri* | Endangered | Stable |
| Birds | *Pterodroma cahow* | Endangered | Increasing |
| Birds | *Pterodroma madeira* | Endangered | Stable |
| Birds | *Rhyticeros narcondami* | Endangered | Stable |
| Birds | *Xenoperdix udzungwensis* | Endangered | Stable |
| Birds | *Amazilia alfaroana* | Critically Endangered | Stable |
| Birds | *Aythya innotata* | Critically Endangered | Stable |
| Birds | *Cyanoramphus malherbi* | Critically Endangered | Stable |
| Birds | *Gymnogyps californianus* | Critically Endangered | Increasing |
| Birds | *Hypotaenidia owstoni* | Critically Endangered | Increasing |
| Birds | *Pterodroma magentae* | Critically Endangered | Increasing |
| Birds | *Strigops habroptila* | Critically Endangered | Increasing |
| Birds | *Vini ultramarina* | Critically Endangered | Stable |
| Bony Fishes | *Centropyge nahackyi* | Near Threatened | Stable |
| Bony Fishes | *Holacanthus limbaughi* | Near Threatened | Stable |
| Bony Fishes | *Holacanthus clarionensis* | Vulnerable | Stable |
| Cone Snails | *Conus kersteni* | Near Threatened | Stable |
| Cone Snails | *Conus fontonae* | Vulnerable | Stable |
| Cone Snails | *Conus teodorae* | Vulnerable | Stable |
| Cone Snails | *Conus xicoi* | Vulnerable | Stable |
| Conifers | *Abies bracteata* | Near Threatened | Stable |
| Conifers | *Abies kawakamii* | Near Threatened | Stable |
| Conifers | *Agathis atropurpurea* | Near Threatened | Stable |
| Conifers | *Agathis microstachya* | Near Threatened | Stable |
| Conifers | *Austrocedrus chilensis* | Near Threatened | Increasing |
| Conifers | *Chamaecyparis lawsoniana* | Near Threatened | Increasing |
| Conifers | *Cryptomeria japonica* | Near Threatened | Stable |
| Conifers | *Halocarpus kirkii* | Near Threatened | Stable |
| Conifers | *Libocedrus bidwillii* | Near Threatened | Stable |
| Conifers | *Libocedrus plumosa* | Near Threatened | Increasing |
| Conifers | *Pherosphaera hookeriana* | Near Threatened | Stable |
| Conifers | *Pinus balfouriana* | Near Threatened | Stable |
| Conifers | *Pinus jaliscana* | Near Threatened | Stable |
| Conifers | *Amentotaxus formosana* | Vulnerable | Stable |
| Conifers | *Callitris oblonga* | Vulnerable | Stable |
| Conifers | *Cupressus macrocarpa* | Vulnerable | Stable |
| Conifers | *Prumnopitys ladei* | Vulnerable | Stable |
| Conifers | *Thuja sutchuenensis* | Endangered | Increasing |
| Conifers | *Juniperus bermudiana* | Critically Endangered | Increasing |
| Cycads | *Encephalartos septentrionalis* | Near Threatened | Stable |
| Cycads | *Macrozamia longispina* | Near Threatened | Stable |
| Cycads | *Zamia pseudomonticola* | Near Threatened | Stable |
| Cycads | *Cycas semota* | Vulnerable | Stable |
| Dicots | *Cleistocactus acanthurus* | Near Threatened | Stable |
| Dicots | *Echinocactus parryi* | Near Threatened | Stable |
| Dicots | *Echinocereus websterianus* | Near Threatened | Stable |
| Dicots | *Mammillaria boolii* | Near Threatened | Stable |
| Dicots | *Pachycereus lepidanthus* | Near Threatened | Stable |
| Dicots | *Parodia columnaris* | Near Threatened | Stable |
| Dicots | *Rebutia arenacea* | Near Threatened | Stable |
| Dicots | *Rhipsalis olivifera* | Near Threatened | Stable |
| Dicots | *Cephalocereus nizandensis* | Vulnerable | Stable |
| Dicots | *Discocactus horstii* | Vulnerable | Stable |
| Dicots | *Gymnocalycium marianae* | Vulnerable | Stable |
| Dicots | *Mammillaria multidigitata* | Vulnerable | Stable |
| Dicots | *Mammillaria tayloriorum* | Vulnerable | Stable |
| Dicots | *Neobuxbaumia polylopha* | Vulnerable | Stable |
| Dicots | *Schlumbergera microsphaerica* | Vulnerable | Stable |
| Dicots | *Cleistocactus sulcifer* | Endangered | Stable |
| Reptiles | *Bradypodion dracomontanum* | Near Threatened | Stable |
| Reptiles | *Furcifer timoni* | Near Threatened | Stable |
| Reptiles | *Kinyongia oxyrhina* | Near Threatened | Stable |

Supplementary figures

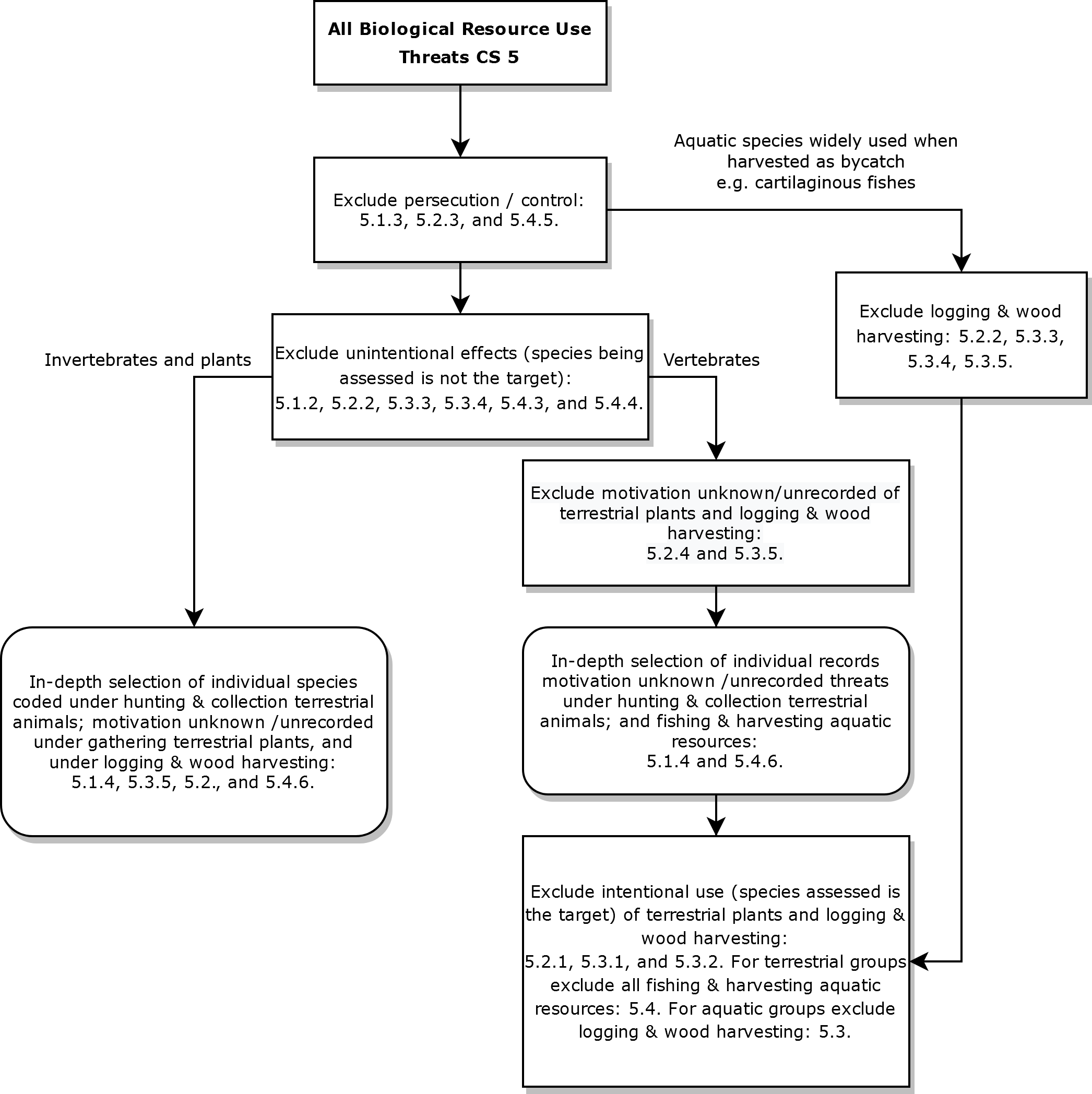

Figure S1. Flow chart describing the process for including forms of biological resource use (Threats Classification Scheme 5) that directly target the species in the taxonomic groups selected, focusing especially on intentional forms of harvest. In-depth selection entails checking a species’ threat categories against detailed text information in the assessment records.

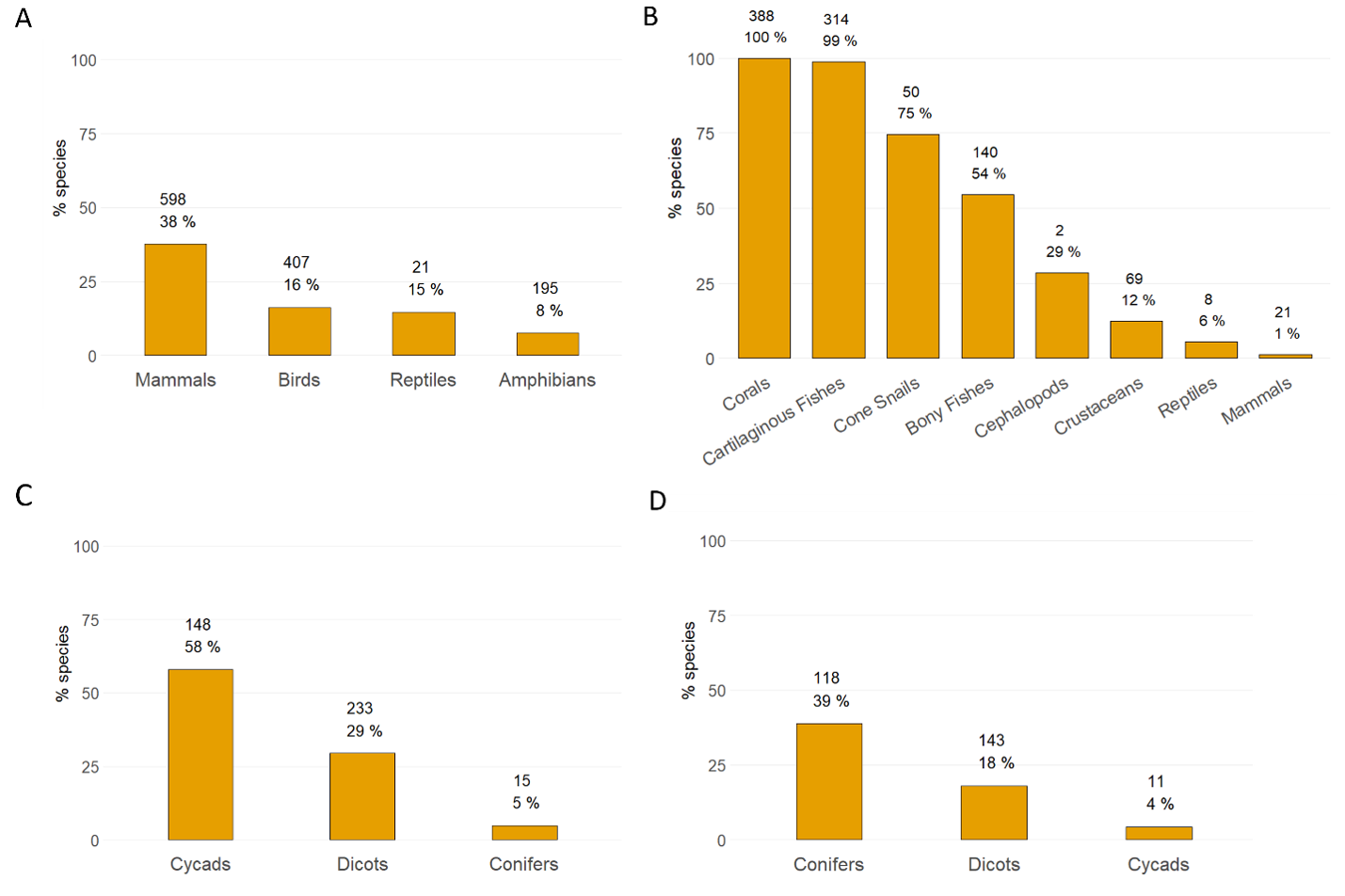

Figure S2. Proportion of NT and threatened species affected by different forms of intentional biological resource use: (A) hunting and collection of terrestrial animals, (B) fishing and harvesting aquatic resources, (C) gathering terrestrial plants, and (D) logging and wood harvesting. The count and percentage of species in the taxonomic group that are affected by the type of use are indicated at the top of bars (minimum estimate only). Some groups contain both terrestrial and aquatic species (e.g. 47 reptile species are affected by hunting and collection of terrestrial animals, and 10 by fishing and harvesting aquatic resources). Note: we reassigned four species that were incorrectly coded on the Red List, namely: *Glyptostrobus pensilis* (CR conifer), incorrectly coded as 5.4.1; *Euastacus brachythorax* (EN, crayfish), incorrectly coded as 5.1.1 (instead of 5.4.1); *Cambarus setosus* (NT, crayfish), incorrectly coded as 5.1.1 (instead of 5.4.1); and *Conus boschorum* (NT, cone snail), incorrectly coded as 5.1.1 (instead of 5.4.1). Bony fishes, dicotyledons (dicots) and reptiles include selected higher-level taxa (Table S1).

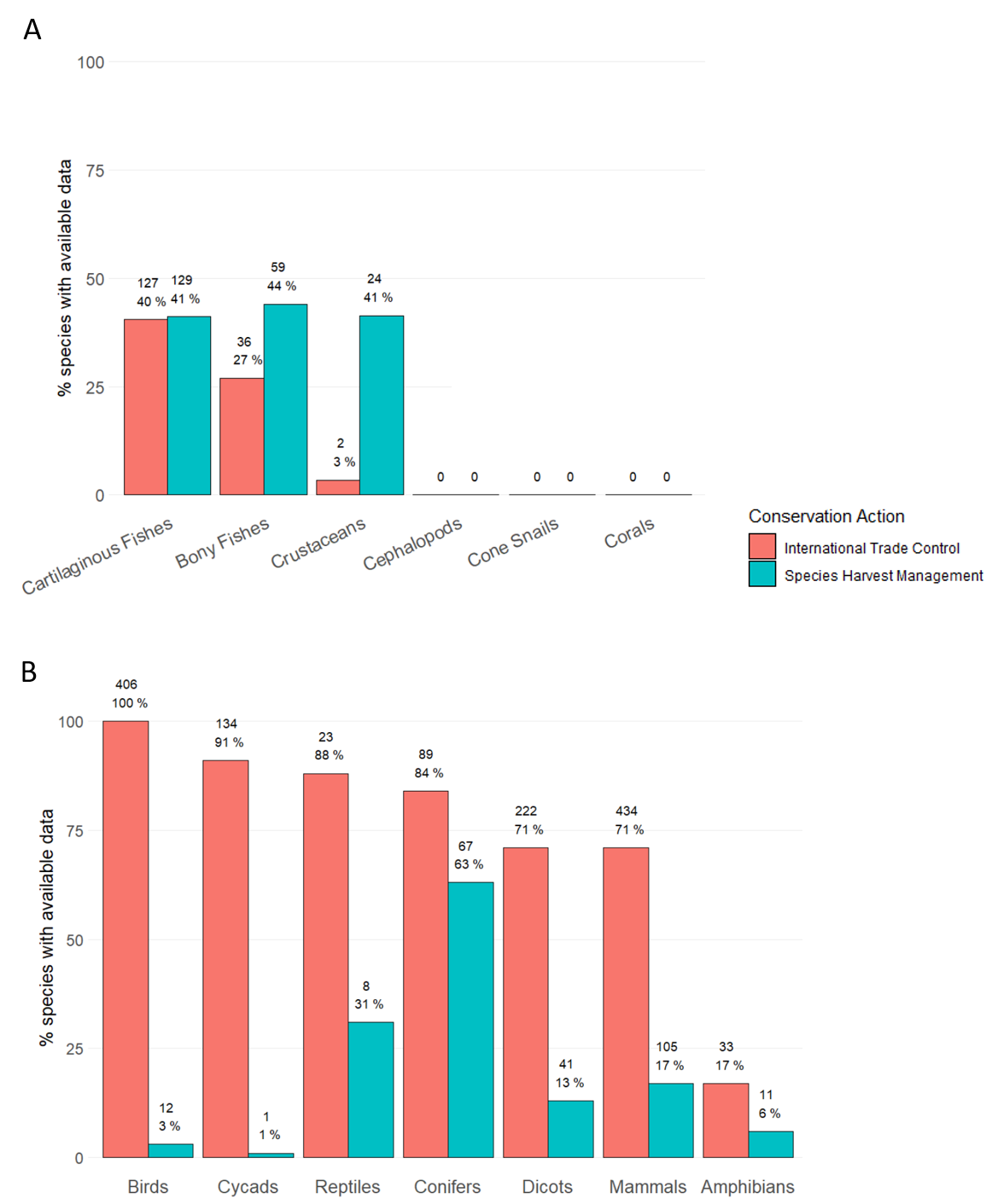

Figure S3. The percentage of NT and threatened species, affected by intentional biological resource use (minimum estimate), with conservation actions data documented for species in (A) aquatic and (B) terrestrial groups. Pink bars (left) denote the percentage of species for which the “International trade control” field is coded as “Unknown,” “Yes” (meaning the species receive such management), or “No” (meaning the species does not receive such management), rather than left blank; blue bars (right) denote the percentage of species for which the “Species Management Harvest Plan” field is coded as “Unknown”, “Yes”, or “No”. Species counts and percentages are labeled in black. Bony fishes, dicotyledons (dicots) and reptiles include selected higher-level taxa (Table S1).

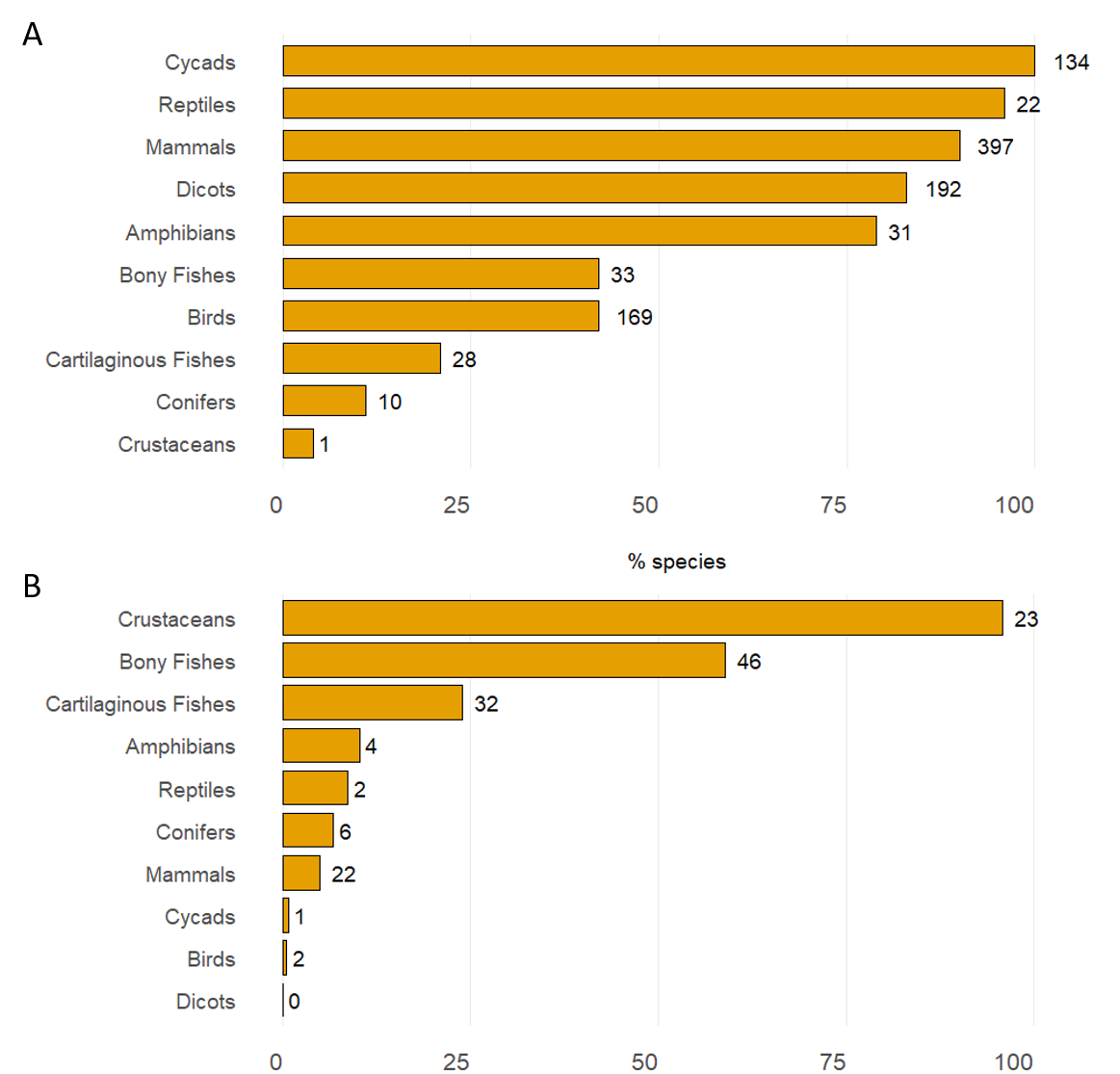
 Figure S4. The percentage of NT and threatened species with available data on conservation actions (those where the field is coded as either “Unknown,” “Yes,” or “No”, rather than left blank) and affected by intentional biological resource use (minimum estimate), receiving (A) international trade control management and (B) targeted species harvest management. No data are available for cephalopods, cone snails or corals (see Figure S3). Bony fishes, dicotyledons (dicots) and reptiles include selected higher-level taxa (Table S1).

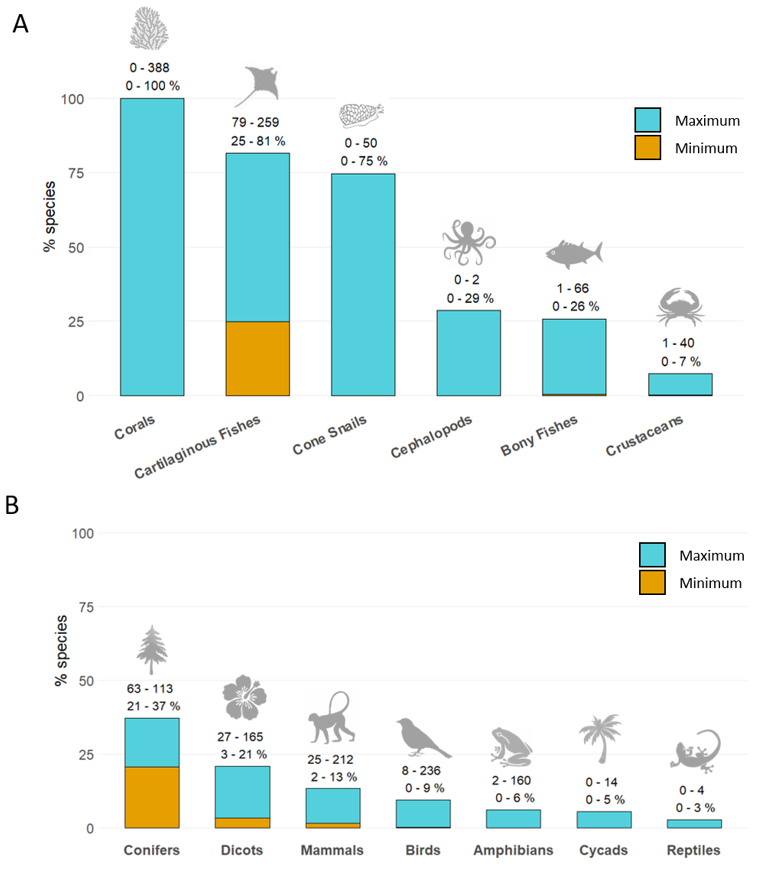

Figure S5. Proportion of NT and threatened species in A) aquatic and B) terrestrial groups, with biological resource use documented as a threat on the Red List and not receiving species conservation management interventions to directly address that use. Minimum (orange) bars show species on the Red List not receiving either international trade control or species harvest management actions (i.e., coded as “No”); maximum (blue) bars include those species where management actions are either not documented for the species (field is left blank), or it is not known whether the species receives those actions (coded as “Unknown”). Labels denote the range of minimum to maximum species counts, and range of minimum to maximum percentages of species within each group. Bony fishes, dicotyledons (dicots) and reptiles include selected higher-level taxa (Table S1).

1. Taxonomic groups qualify for inclusion according to criterion i if they have >40% of LC, threatened, and all extant, data sufficient species with at least one purpose of use coded; taxonomic groups meet criterion ii when the proportion of LC species with at least one purpose of use code falls above or within the range of the proportion of species with Use and Trade coding across the other Red List categories.
2. We excluded past threats which were deemed unlikely to return. Where severity or scope of threat is coded as unknown, we assign each a score of 2 (medium), meaning threats whose severity or scope are unknown are still analyzed if their timing is coded as future or likely to return. Consequently, for threats where both severity and scope are coded as unknown, our approach is precautionary in assuming the threat is at least medium impact; for threats where one of severity or scope are coded as unknown, and the other as either slow/fluctuating or negligible / low impact, our approach is evidentiary in assuming that the threat is not medium impact.
